# RNA polymerase I transcription fidelity, speed and processivity depend on the interplay of its lobe binding subunits

**DOI:** 10.1101/433375

**Authors:** Philipp E. Merkl, Michael Pilsl, Tobias Fremter, Gernot Längst, Philipp Milkereit, Joachim Griesenbeck, Herbert Tschochner

## Abstract

Eukaryotic RNA polymerases I and III (Pol I and III) consist of core subunits, which are conserved in RNA polymerase II (Pol II). Additionally, Pol I and III have specific subunits, associating with the so-called ‘lobe’ structure first described within Pol II. In Pol I of the yeast *S. cerevisiae*, these are Rpa34.5, and the N-terminal domains of Rpa49 and Rpa12.2, here referred to as the lobe-binding module (lb-module). We analyzed functions of the lb-module in a defined *in vitro* transcription system. Cooperation between lb-module components influenced transcription fidelity, elongation speed, and release of stalled Pol I complexes to continue elongation. Interestingly, lb-module containing Pol I and III, but not Pol II, were able to transcribe nucleosomal templates. Our data suggest, how the Pol I specific subunits may contribute to accurate and processive transcription of ribosomal RNA genes.

## Introduction

The eukaryotic RNA polymerases I, II and III (Pol I, II, III) have a similar architecture and contain ten conserved subunits which form the catalytic core. Furthermore, they share a more distantly related heterodimer at the periphery of the enzyme, designated as the stalk, formed by Rpa14/Rpa43, Rpb4/Rpb7 and Rpc17/Rpc25 in Pol I, Pol II and Pol III, respectively (see as review (Khatter et al., 2017)). In the yeast *S. cerevisiae* Pol I and Pol III have additional subunits associated. The heterodimer Rpa34.5/Rpa49 of Pol I corresponds to Rpc37/Rpc53 of Pol III (Khatter et al., 2017) and associates together with the N-terminal part of Rpa12.2 to the Pol I lobe structure forming a lobe binding module (lb-module). Based on structural homology, localisation on the core enzyme and function Rpa34.5/Rpa49 are considered as counterparts of the Pol II transcription initiation factors TFIIE and TFIIF (Kuhn et al., 2007, Jennebach et al., 2012, Geiger et al., 2010). The heterodimer Rpa34.5/Rpa49 consists of a dimerization module formed by Rpa34.5 and the N-terminal part of Rpa49 (full length Rpa34.5 and aa 1–110 of Rpa49), the A49 linker (aa 105–187 of Rpa49) and the C-terminal part of Rpa49 (Rpa49CT, aa 187–415). The dimerization module binds to the Pol I lobe domain on the side of the active center which is opposite to the stalk (Jennebach et al., 2012, Fernandez-Tornero et al., 2013, Engel et al., 2013) and occupies a similar position as the TFIIF dimerization module in association with Pol II (yeast Tfg1/Tfg2) (Geiger et al., 2010, Kuhn et al., 2007). The C-terminal part of Rpa49 forms a tandem winged helix (tWH) similar as in the small TFIIE subunit Tfa2 (Geiger et al., 2010) (Okuda et al., 2000) or in RPC37 (Hoffmann et al., 2015). Although this domain cannot be unambiguously localized in current structures of Pol I, there is evidence for a preferential localization at the DNA binding cleft of Pol I at the interface between the stalk and the upstream DNA (Jennebach et al., 2012, Pilsl et al., 2016b, Engel et al., 2013, Tafur et al., 2016).

Interplay between the Pol II factors TFIIF and the RNA cleavage factor TFIIS was reported to promote transcript elongation and increase transcription through nucleosomes in a synergistic manner (Schweikhard et al., 2014, Luse et al., 2011). Similar to the Pol II counterparts the Pol I subcomplex Rpa34.5/Rpa49 was suggested to enhance RNA elongation and cleavage (Kuhn et al., 2007, Liljelund et al., 1992, Beckouet et al., 2008, Geiger et al., 2010) and to stabilize the initiation complex (Tafur et al., 2016, Jennebach et al., 2012), presumably contributing to a high rate of Pol I loading (Albert et al., 2011). Interestingly, polymerase-dependent RNA cleavage is stimulated by the Rpa34.5/Rpa49 dimerization module (Geiger et al., 2010). This is important to extend arrested and backtracked elongation complexes by removal of a displaced RNA 3’end from the active center of the enzyme (Lisica et al., 2016). The stimulatory effect was explained as an indirect effect resulting from its proximity to subunit Rpa12.2 (Jennebach et al., 2012), whose C-terminal part is required for cleavage (Kuhn et al., 2007). Rpa49CT binds DNA and supports both promoter-dependent transcription and extension of RNA from a transcription bubble (Pilsl et al., 2016a, Geiger et al., 2010). Based on its position on the Pol I enzyme it was speculated to assist promoter escape by keeping the Pol I clamp structure closed and stabilizing the Pol I- interaction with DNA upstream of the start site (Sadian et al., 2017, Tafur et al., 2016). *In vivo*, expression of Rpa49CT, but not of Rpa34.5/Rpa49NT is required for normal cell growth as well as for recruitment of Pol I to the rRNA gene (Beckouet et al., 2008).

The Pol I lb-module contains not only the dimerization module Rpa34.5/A49NT but also the N-terminal part (Rpa12ΔC, aa 1- aa 85) of subunit Rpa12.2. Rpa12.2 is homologous to the Pol II transcription elongation factor TFIIS. Deletion of Rpa12.2 results in growth inhibition at elevated temperature, sensitivity to nucleotide-reducing drugs, inefficient transcription termination and hampers the assembly of the Pol I enzyme (Nogi et al., 1993, Van Mullem et al., 2002, Prescott et al., 2004). The lack of the complete Rpa12.2 may also lead to the loss of Rpa34.5/Rpa49, as judged by immunoprecipitation of Pol I from a *rpa12Δ* strain (Van Mullem et al., 2002). If only the C-terminal part of Rpa12.2 is lacking, Rpa34.5/Rpa49 remain Pol I-associated. Whereas, deletion of the C-terminal Rpa12.2 domain showed no significant growth phenotype under the tested conditions, deletion of the N-terminus led to growth phenotypes indistinguishable from that of a *rpa12Δ* strain (Van Mullem et al., 2002). The lack of a growth phenotype of an *rpa*12ΔC strain was surprising, since it is important for RNA cleavage activity when transcription is stalled (Kuhn et al., 2007, Lisica et al., 2016) (Van Mullem et al., 2002). Recent structural investigations indicated that the C-terminal part of Rpa12.2 is displaced from the active site of the enzyme when Pol I is bound to the transcription initiation factor Rrn3 (Pilsl et al., 2016a, Engel et al., 2016, Tafur et al., 2016). Furthermore, it was proposed that Rpa12.2 is an intrinsic destabilizer of elongating Pol I (Appling et al., 2018) and might release - at least under specific conditions - Rpa34.5/Rpa49 from the lobe (Tafur et al., 2018). In summary, it can be concluded that Rpa12.2 in addition to Rpa34.5/Rpa49 is important for the elongating Pol I. The lobe associated N-terminal domain of Rpa12.2 is thereby crucial for Pol I assembly, whereas the C-terminal domain is dynamic and supports RNA cleavage.

Here, we investigated the function of the Rpa34.5/49 heterodimer and its interplay with Rpa12.2 using defined promoter-dependent and -independent *in vitro* transcription systems. In particular our studies emphasize the roles of the dimerization domain and Rpa49CT for transcription fidelity, Pol I speed and resumption of stalled Pol I complexes. We also tested the possibility that these subunits and Rpa12.2 contribute to transcription of nucleosomal templates *in vitro* and compared the three nuclear RNA polymerases in their efficiency to transcribe such templates.

## Results

### Domains of the Pol I-specific heterodimer Rpa34.5/Rpa49 influence elongation rate and Pol I processivity

To study the influence of Pol I lb-module subunits on *in vitro* transcription wildtype and mutant Pol I were affinity-purified through the ProtA-tagged subunit Rpa135 using the same stringent salt conditions (Merkl et al., 2014). Purified WT Pol I contained all 14 subunits, ΔRpa49 Pol I lacked the heterodimeric subunit Rpa34.5/Rpa49, ΔRpa12 Pol I lacked subunit Rpa12.2 and the heterodimer Rpa34.5/Rpa49, and Rpa12ΔC Pol I contained the N-terminal part of Rpa12.2 and the Rpa34.5/Rpa49 heterodimer (Suppl Fig. 1A, B), in good correlation with previous analyses (Van Mullem et al., 2002). To analyse the effects of Rpa34.5/Rpa49 or its subdomains on *in vitro* transcription, the recombinant heterodimer or protein fragments of the truncated heterodimer were purified (Suppl. Fig. 1C) and added to affinity purified WT Pol I or Pol I depleted in lb-module forming subunits.

The activity of wildtype Pol I and Pol I mutants in elongation *in vitro* was previously analysed using various DNA/RNA hybrid scaffolds (Kuhn et al., 2007, Geiger et al., 2010) which might differ from transcription bubbles generated by promoter-dependent Pol I transcription. Therefore, we compared transcription activity of different Pol I enzymes in promoter-dependent elongation assays using templates with the Pol I promoter followed by 169nt of DNA lacking guanines in the sense strand (G-less cassette) which were immobilized at the 3’ end on magnetic beads (Fig. 1 A). Using this template transcription could be specifically initiated at the promoter in a minimal system consisting of Pol I and recombinant Rrn3 and CF. If the transcription buffer contained only UTP, α^32^P-CTP, and ATP, but no GTP, ternary complexed were stalled at the end of the G-less cassette (Fig. 1B, lanes 1, 6). Ternary arrested complexes were washed with buffers containing ATP, CTP and UTP to remove unincorporated ^32^P-CTP and unbound proteins. After addition of GTP (chase), time course experiments were performed to follow transcript extension (Fig. 1B/C). The amounts of fully extended transcripts divided by the amount of arrested transcripts at time point zero was used as a measure for Pol I processivity. Using WT Pol I more than 90% of stalled transcripts could be extended and reached the full-size product. In contrast, using a ΔRpa49 Pol I only around 60% of the arrested transcripts were fully extended after 2 minutes chase indicating reduced Pol I processivity. The escape of ΔRpa49 Pol I from the arrest after the G-less cassette was apparently not impaired, since for both, WT and ΔRpa49 Pol I, we observed that less than 12% of the transcripts could not be further elongated after NTP addition. It is thus likely, that ΔRpa49 processivity was affected downstream of the G-less cassette after resuming elongation. Looking at the migration pattern of radiolabelled RNAs at early timepoints indicated a slightly slower elongation speed of ΔRpa49 Pol I if elongation was performed at room temperature in the presence of 200 μΜ NTPs (see arrows in Fig. 1B)(see text to Fig. 3).

**Fig. 1.**
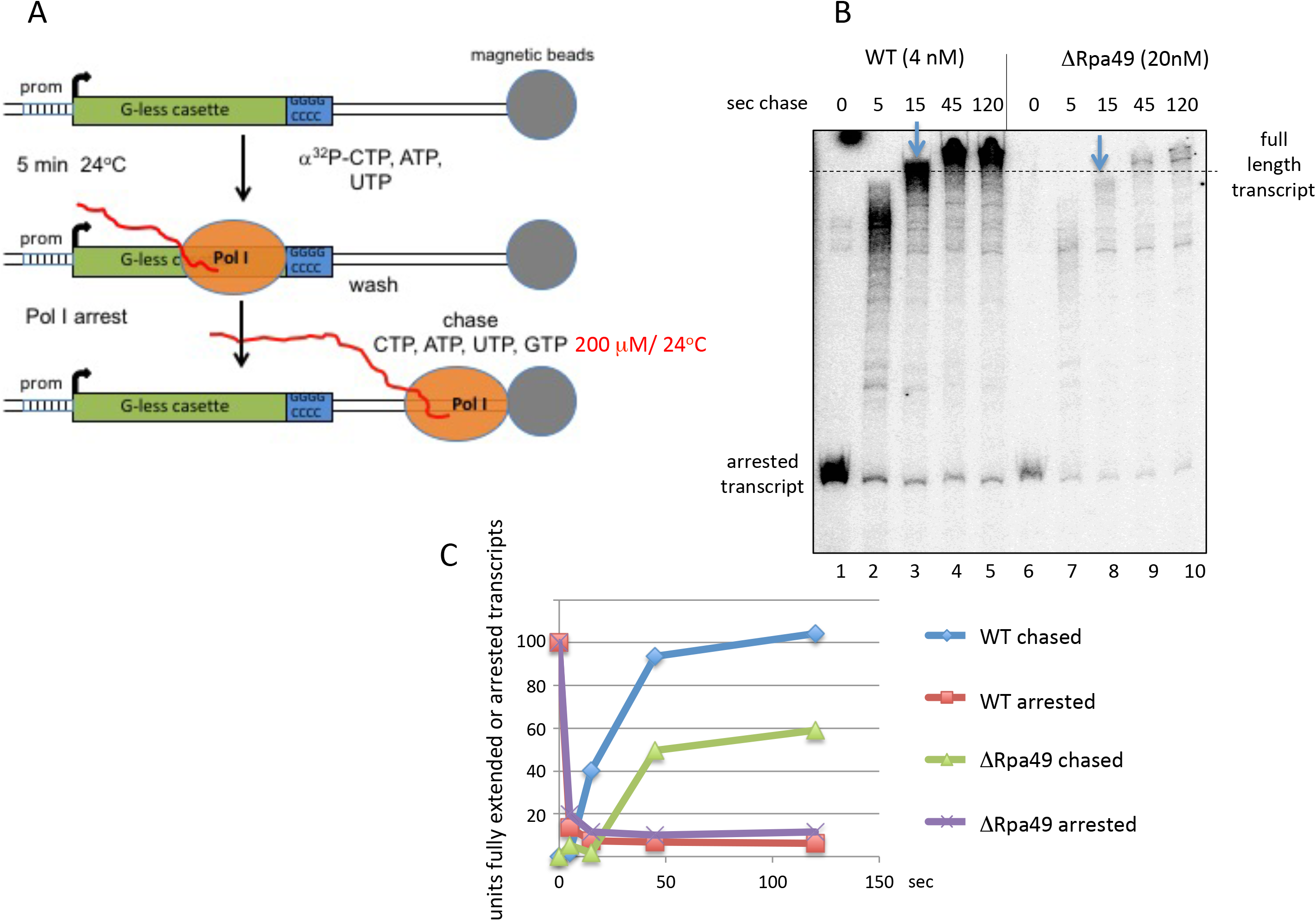
Depletion of the heterodimeric Rpa34.5/49 complex results in reduced Pol I processivity. A) Experimental design. Promoter-dependent transcription was performed for 10min on templates attached to magnetic beads using 70nM Rrn3, 20nM CF and 4nM Pol I and in the presence of 200 μM ATP, 200 μM UTP, 10 μM CTP (^32^P-CTP) but no GTP. Transcription was arrested after 169 nt when the first GTP should be incorporated. Arrested ternary complexes were washed twice with excess transcription buffer without NTPs and radioactivity and were chased adding 200 μM NTPs including GTP. At the time points indicated 25 μl aliquots were withdrawn and injected in 200 μl 0.6% SDS, 0.3 M NaCl, 5 mM EDTA, 0.5 mg/ml Proteinase K, which immediately stopped rRNA synthesis. B) Promoter-dependent elongation assays were performed on template PIP_G-extend 2kb which produces a 1759 nt long run off transcript after the elongation block using 4 nM WT or 20 nM ΔRpa49 Pol I. Pulse labelling was for 5 min. Chase conditions were 200 μM NTPs and 24°C. Radiolabelled transcripts were separated on a denaturing polyacrylamide/urea gel and detected using a PhosphorImager. The arrows indicate that at time point 15’ RNA extension of the WT enzyme was slightly advanced. C) Comparative analysis of Pol I processivity between WT and ΔRpa49 enzyme of the experiment shown in (B). Percentage of fully extended and arrested transcripts in reference to the pulse labelled arrested transcript (time point 0) at the respective time points. (blue and green: fully extended transcripts of WT and ΔRpa49; orange and purple: arrested transcript of WT and ΔRpa49, respectively). The experiment is representative for multiple replicates n>4.

### Pol I mutants lacking the lobe-binding-module or the C-terminal part of Rpa12.2 impair Pol I processivity in distinct ways

To see how components of the Pol I lb-module influence Pol I processivity, ternary complexes of WT, ΔRpa12, Rpa12ΔC and ΔRpa49 substituted with Rpa49CT were arrested at the end of the G-less cassette for 20 min. Addition of 200 μΜ NTPs to the washed ternary complexes led to full extension of less than 50% of the arrested transcripts at the G-less cassette in the three Pol I mutants tested, whereas most transcripts could be completely extended by WT Pol I (Fig 2A and B). This indicated that Pol I processivity was impaired when the integrity of the lb-module was affected (ΔRpa12 ΔRpa49 + Rpa49CT), or the C-terminal part of Rpa12.2 was missing (Fig, 2 A and B). When elongation was performed with reduced NTP concentration (20 μΜ NTPs) Pol I processivity was reduced in all Pol I preparations (Suppl. Fig. 2). However, whereas in WT Pol I still almost 60% of the arrested transcripts could be fully extended, processivity in mutant Pol I enzymes was even more impaired. After 60 second of chase not more than 30 % of the arrested transcripts reached the full size. This underlines the importance of the lb-module, in particular the dimerization domain Rpa34.5/Rpa49NT and subunit Rpa12.2, for efficient elongation. Interestingly, the way how processivity was affected differed for the mutant enzymes. In the presence of Rpa12ΔC only about 40% transcripts stalled at the G-less cassette could complete chain elongation after 60 seconds chase, whereas more than 85% of the arrested transcripts could be extended, when WT, ΔRpa12.2, and ΔRpa49 + Rpa49CT polymerases resumed chain elongation (Fig. 2C and Suppl. Fig. 2). This could mean that in Rpa12ΔC Pol I, which contains Rpa34.5/Rpa49, cleavage is required for efficient restart of transcription. In contrast ΔRpa12.2 which is also impaired in cleavage but lacks Rpa34.5/Rpa49 was more efficient in the elongation of arrested transcripts. Rather efficient resumption of elongation could be also observed for ΔRpa49 + Rpa49CT Pol I which contained Rpa12.2 but no dimerization module. Accordingly, it is possible that the dimerization module keeps Pol I in an arrested and backtracked state until the displaced 3’end of the transcript can be cleaved. Consistently, in the absence of the dimerization module resumption of elongation is possible, no matter if Rpa12.2 is present or not. On the other hand, ΔRpa12.2 and ΔRpa49 mutants were able to resume elongation of most arrested transcripts at the G-less cassette but could not complete full extension. This could be due to structural challenges when transcription of a long template leads to torsions and/or the generation of long DNA-RNA hybrids interfering with the elongating enzyme. Another interesting finding in this set of experiments was that the mutant polymerases Rpa12.2 AC, and ΔRpa49 + Rpa49CT did not stop RNA synthesis precisely at the G-less cassette, suggesting that wrong nucleotides were incorporated (Fig. 2A, see also results below and Fig. 4).

**Fig. 2.**
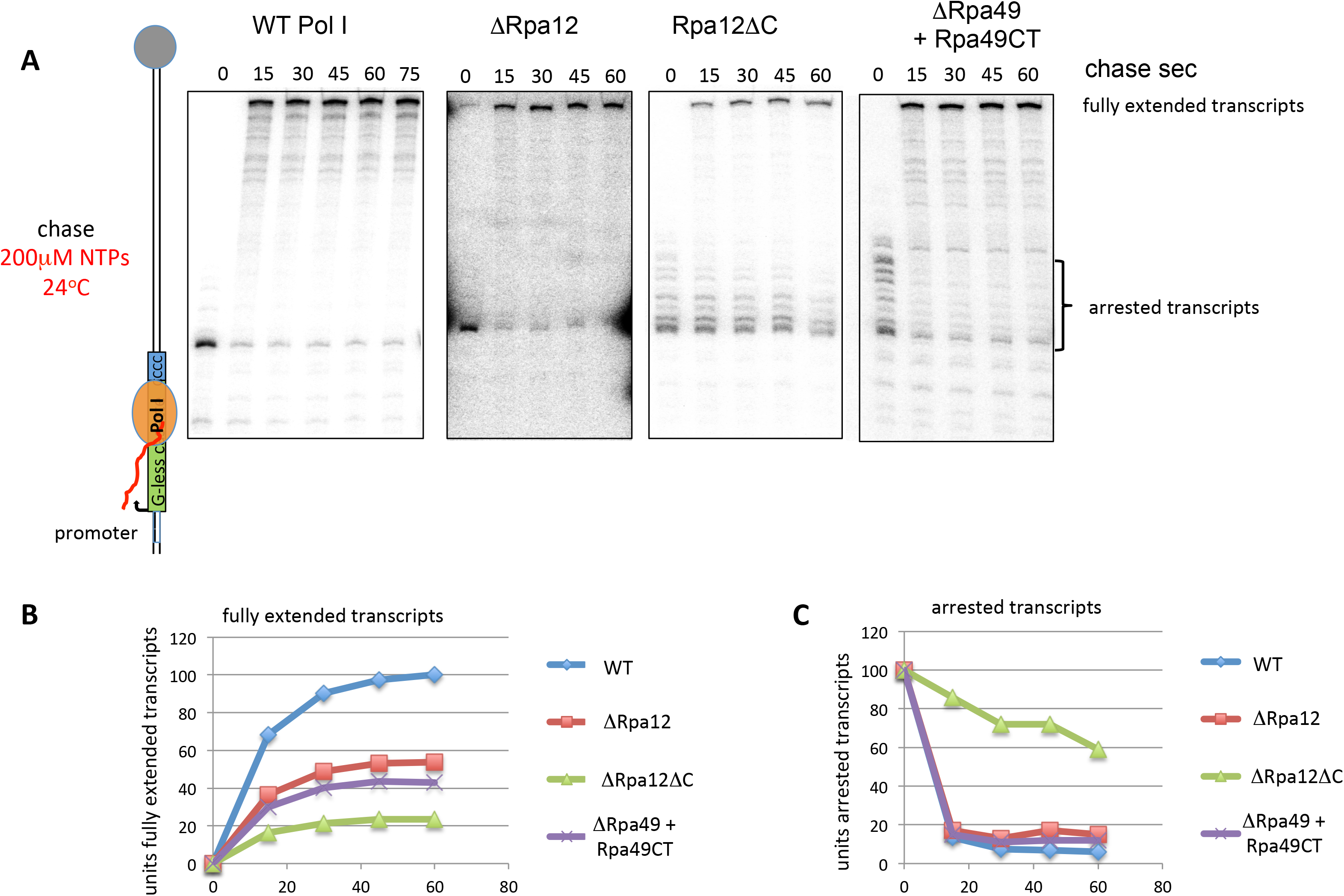
Reduction of Pol I processivity appears in different mutant enzymes when arrested transcripts should be extended at a G-less cassette. A) Promoter-dependent elongation assays were performed as described in Fig. 1, with the exception that the initial pulse was performed for 20 min and that the separation distance of the transcripts on the gel was longer. Assays were performed using 4nM Pol I RpaA12 and ΔRpa49 and 12 nM ΔRpa12. Each experiment is representative for at least three replicates (see also Suppl Fig 2.). B) Percentage of fully extended transcripts in reference to the pulse labelled arrested transcript (time point 0) at the respective time points. C) Percentage of arrested transcripts in reference to the pulse labelled arrested transcript (time point 0) at the respective time points.

### The tWH of Rpa49CT accelerates Pol I speed

The migration pattern of radiolabelled transcripts at early timepoints after the NTP chase indicated a slightly slower elongation speed of ΔRpa49 Pol I than WT Pol I if elongation was conducted at room temperature in the presence of 200 μM NTPs (see arrows in Fig. 1B). Furthermore, addition of Rpa49CT to ΔRpa49 resulted in an apparent faster Pol I movement compared to WT Pol I when elongation was performed in the presence of 20 μM NTPs (Suppl. Fig. 2, compare time points at 30 sec). Therefore, we studied the impact of the Rpa34.5/Rpa49 domains on elongation speed in more detail. When *in vitro* promoter-dependent transcription reactions were carried out at lower temperature, the influence of the Rpa49CT and Rpa34.5/Rpa49NT domains on the transcription rate was better detectable (Fig. 3A). As previously described (Pilsl et al., 2016a), addition of either the heterodimer Rpa34.5/Rpa49 or Rpa49CT to the initiation inactive ΔRpa49 Pol I supported transcription initiation (Fig. 3 A, compare lanes 9–12 and 17 to 20 with 13 to 16). Addition of the heterodimer Rpa34.5/Rpa49 resulted in a similar elongation speed as WT Pol I (Fig. 3A, compare lanes 1–4 with 5–8/ Suppl. Fig. 3). Interestingly, addition of Rpa49CT increased the elongation speed in promoter-dependent transcription at 0°C, (Fig. 3A, compare lanes 17–21 with 9–12). Elongation speed of ΔRpa49 Pol I was also investigated in promoter-independent transcription assays using tailed templates. In these reactions the presence of Rpa49CT also resulted in increased elongation speed in the presence of 200μM NTPs, when compared with WT polymerase. In contrast, addition of the Rpa34.5/Rpa49NT dimerization domain did not significantly change elongation speed of ΔRpa49 Pol I (Fig. 3B). These experiments indicate that the complete heterodimer is necessary to restore WT processivity of ΔRpa49 Pol I, and that the presence of Rpa49CT leads to an increased elongation speed.

**Fig. 3.**
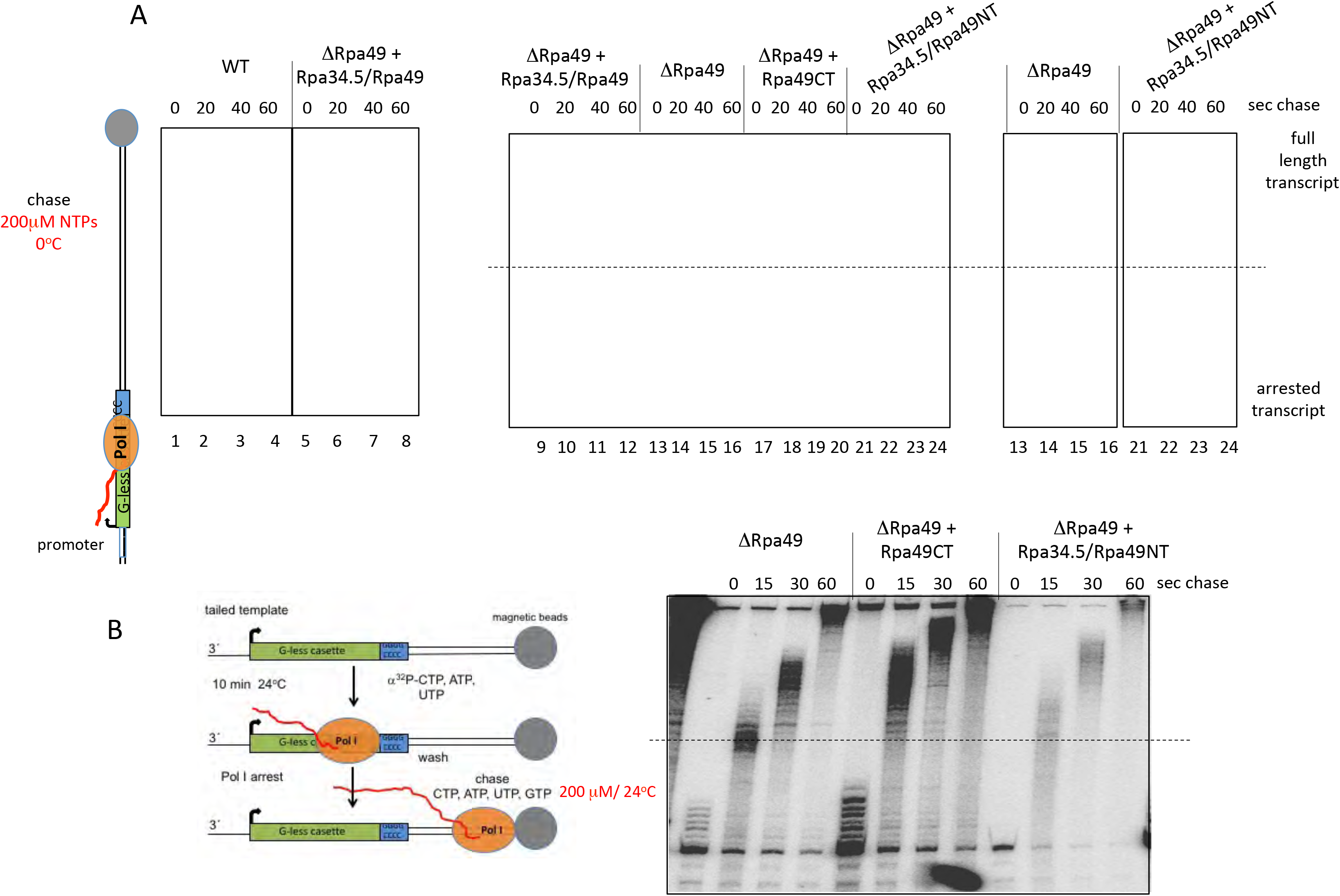
Pol I containing Rpa49CT but lacking the dimerization module Rpa34.5/49NT accelerates transcription. A) Promoter-dependent elongation assays were performed as described in Fig. 1. Combinations of different Rpa34.5/Rpa49 fractions were added to ΔRpa49 (4nM) before transcription was started. Either 50 nM purified recombinant heterodimer Rpa34.5/Rpa49 was used or the dimerization domain Rpa34.5/Rpa49NT lacking the aminoacids 181 to 426 of Rpa49 or the C-terminal part of Rpa49 (Rpa49CT) which corresponds to a protein fragment from aminoacid 111 to 426 of Rpa49. Pulse labelling was for 5 min. Under this rather short labelling conditions most transcripts stopped at the end of the G-less cassette. Chase conditions were 200 μM NTPs and 0°C. Radiolabelled transcripts were separated on a denaturing polyacrylamide/urea gel and detected using a PhosphorImager. Elongation was faster in the presence of Rpa49CT. The original gel section is shown in Suppl. Fig. 3. Longer exposed gel sections of lanes 13-16 and 17–20 are shown on the right. Experiments were repeated at least twice (see also Fig. 3B). B) Tailed template transcription to analyse effects of the dimerization domain and Rpa49CT on transcription elongation. Left panel, experimental design was similar as in Fig. 1 with the exception that transcription was initiated at a 3’ overhang of the immobilized template which allowed RNA synthesis without addition of initiation factors. Rpa34.5/Rpa49 or Rpa49CT (each 50 nM) were added to ΔRpa49 and analysed as in A. Elongation was stopped after 15, 30 and 60 seconds, respectively.

### Wrong nucleotides can be incorporated in the nascent RNA chain if the Rpa34.5/Rpa49NT dimerization domain or the C-terminal part of Rpa12.2 is absent

The pattern of radiolabelled transcripts observed before chase in tailed template reactions with ΔRpa49 Pol I, indicated that the enzyme was not efficiently arrested after the G-less cassette (Fig. 3B, first lane). This could be due to misincorporation of NTPs. To investigate a general role of the Rpa49/Rpa34.5 heterodimer in controlling the correct incorporation of nucleotides, we analysed how the Rpa34/Rpa49 subdomains influenced transcription fidelity at the G-less cassette using promoter dependent transcription. WT Pol I and ΔRpa49 Pol I in the presence of Rpa34/Rpa49 domains were used for promoter-dependent *in vitro* transcription using templates with G-less cassettes in the absence of GTP. The reaction was stopped after 5, 15 and 30 minutes and supernatants (su) were separated from the immobilized template (p) to detect possible differences in the stability of the arrested ternary complexes. Whereas WT Pol I and ΔRpa49 Pol I in the presence of Rpa34.5/Rpa49 stopped precisely at the end of the G-less cassette (Fig. 4A lanes 1-2 and lanes 3 - 6), ΔRpa49 Pol I together with Rpa49CT significantly extended the stalled transcript in the absence of GTP (Fig. 4A, compare lanes 7–12 with lanes 1–6). Addition of the purified dimerization module to ΔRpa49 and Rpa49CT polymerases reduced extension of the transcripts (Fig. 4A, lanes 13 - 18), arguing that the presence of Rpa34.5/Rpa49NT prevents elongation of the arrested transcripts. The absence of the dimerization module did not significantly impair the stability of the arrested ternary complex, since the ratio of immobilized transcripts versus transcripts in the supernatant (su) was comparable in all reactions (Fig. 4A, lower panel). Furthermore, washing the arrested complexes with increasing salt concentrations (up to 1 M NaCl) did not result in disassembly of the complexes (data not shown). This suggests that stalled complexes are stable, even in the absence of the dimerization module.

**Fig. 4.**
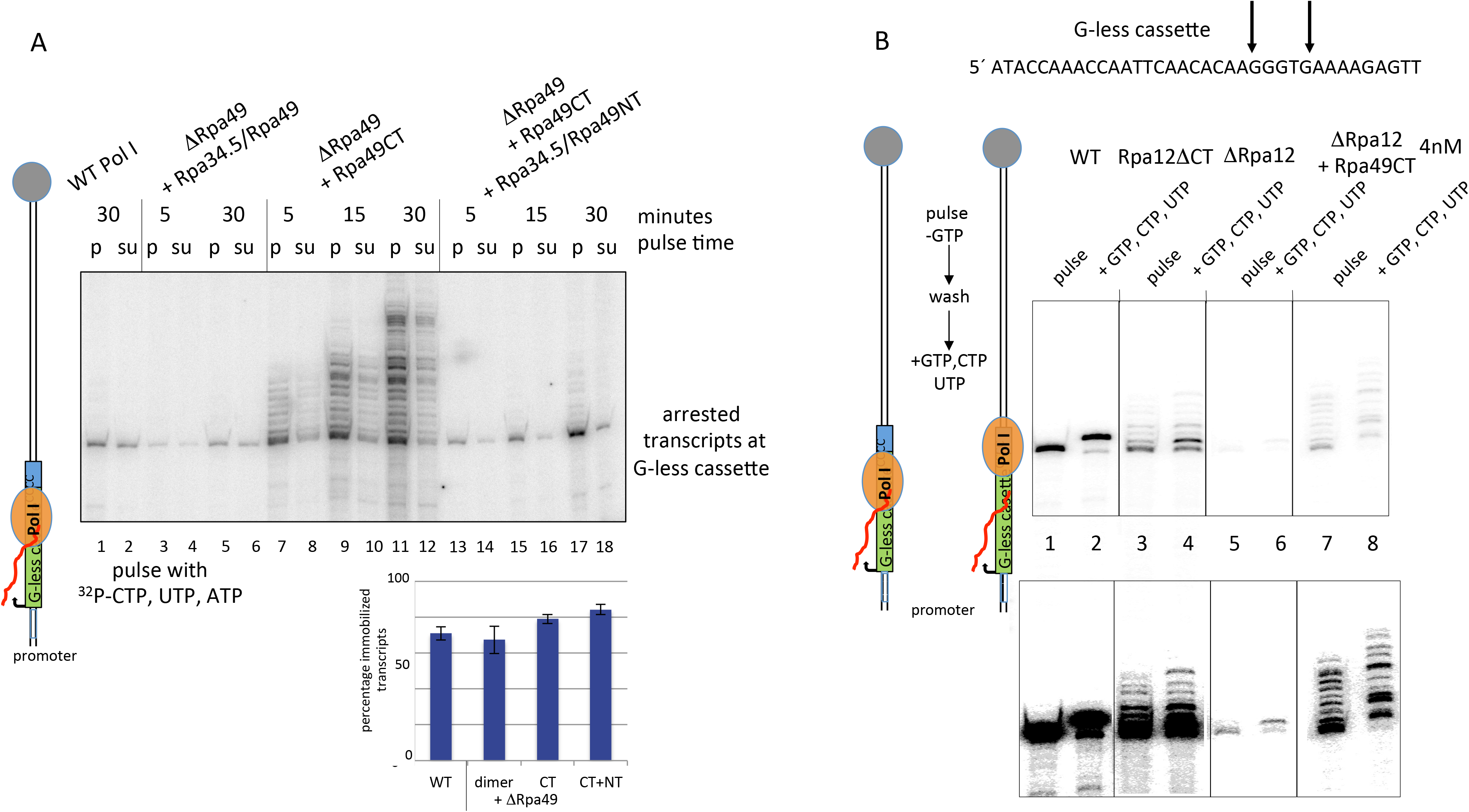
The heterodimer Rpa34.5/Rpa49 and the C-terminal part of Rpa12 are both required for correct incorporation of nucleotides, but influence re-extension of arrested transcripts differently. A) The dimerization module of Rpa34.5/Rpa49 is required to prevent Pol I from mis-incorporation of nucleotides when transcription is arrested at the G-less cassette. *Upper panel* Promoter-dependent transcription assays on immobilized templates containing a G-less casette were performed in the presence of ATP, UTP and α^32^P-CTP using the indicated Pol I fractions (5 nM). After the indicated time points supernatants (su) were separated from pellets (p) and analyzed on a 6% polyacrylamide gel containing 6 M urea. *Lower panel* Ternary stalled transcription complexes are rather stable. Quantitation of the ratio pellet/supernatant of arrested transcripts synthesized on immobilized templates from four different experiments. (dimer: Rpa34.5/Rpa49; CT: Rpa49CT; CT + NT: Rpa49CT + Rpa34.5/Rpa49NT). (Experiment is representative for two replicates). B) Pulse labelling and extension of stalled transcripts of Pol I mutants lacking either the C-terminal part of Rpa12.2 (Rpa12ΔCT), or the entire Rpa12 and Rpa34.5/Rpa49 (ΔRpa12), or Rpa12.2 and the dimerization module (ΔRpa12 + Rpa49CT). Promoter-dependent transcription assays on immobilized templates containing a G-less cassette were performed as in Fig. 1. After washing the ternary complexes with a buffer containing ATP, UTP and CTP (end concentration 200 μM), an aliquot was removed. GTP, UTP and CTP, but no ATP were added to the remaining fraction. Transcription elongation was stopped after 3 min. The lower panel shows a longer exposure of the same gel to detect ΔRpa12-derived transcripts. Transcripts were analyzed on a 6% polyacrylamide gel containing 6 M urea. The DNA sequence of the end of the G-less cassette is indicated at the top. Arrows point to the first GTP and the first ATP after the G-less cassette. (Experiment is representative for two replicates).

Next, we wanted to see whether the Rpa49CT domain can also promote nucleotide misincorporation of ΔRpa12.2 Pol I, lacking Rpa12.2 as well as the Rpa34.5/Rpa49 heterodimer. Therefore, we compared elongation of arrested transcripts in reactions containing, WT Pol I, Rpa12.2ACT Pol I, ΔRpa12.2 Pol I and ΔRpa12.2 Pol I in the presence of Rpa49CT. Addition of Rpa49CT to ΔRpa12.2 Pol I supported RNA synthesis in promoter-dependent transcription (Fig. 4B, compare lane 7 with lane 5) underlining the important role of this domain in initiation (Beckouet et al., 2008, Pilsl et al., 2016a). Furthermore, in contrast to ΔRpa12.2 Pol I, an increased number of readthrough elongated transcripts after the G-less cassette were observed suggesting that the Rpa49CT tWH in the absence of the Rpa34.5/Rpa49NT dimerization domain promoted misincorporation of nucleotides. Interestingly, readthrough elongated transcripts were also detected using Rpa12.2ACT Pol I in transcription reactions (Fig. 4B compare lanes 3 and 1), which suggested that the C-terminus of Rpa12.2 is also required to prevent NTP misincorporation. This is in accordance with a recently published finding (Sanz-Murillo et al., 2018). Apparently, wrong nucleotides can be incorporated by different Pol I mutants, but only in the presence of Rpa12.2 removal of misincorporated nucleotides can occur.

It was recently reported that Rpa12.2AC Pol I is not able to extend arrested and backtracked transcripts which result in a displaced RNA 3’ end from the active enzyme (Lisica et al., 2016). Accordingly, arrested and backtracked transcripts might accumulate after the G-less cassette in such mutants. Extension of arrested and backtracked transcripts should require an RNA 3’ cleavage activity which was reported to depend on the C-terminal domain of subunit Rpa12.2 (aa 79 to aa 125) (Kuhn et al., 2007, Lisica et al., 2016) and on the dimerization module of Rpa34/Rpa49 (Geiger et al., 2010). In fact, if CTP, UTP and GTP, but no ATP were provided for chain elongation after washing the arrested ternary complexes, Rpa12.2AC Pol I could only poorly extend the nascent RNA, while misincorporating nucleotides (Fig. 4B, compare lane 4 with lane 3). In contrast, WT Pol I could further extend most of the radiolabelled RNAs until the position where the first ATP should be incorporated (Figure 4B, compare lanes 1 with 2, see sequence of the sense strand of the DNA template on the top). Interestingly, most transcripts arrested after the G-less cassette could also be extended if Rpa49CT was added to ΔRpa12 Pol I underlining that the presence of the dimerization domain is important to keep the elongating polymerase in the arrested state if the cleavage supporting activity of Rpa12.2 is lacking. On the other hand, the Rpa12ΔC mutant which is lacking the RNA cleavage supporting activity, but contains Rpa34.5/Rpa49 and the N-terminus of Rpa12.2 could not resume efficient chain elongation. It was previously reported that stalled Pol II transcripts due to the absence of one nucleotide can be extended with or without RNA cleavage activity (Izban and Luse, 1993). Such elongation competent ternary complexes which apparently produce mainly shallow backtracks (Izban and Luse, 1993) (Lisica et al., 2016) could also arise in Pol I-dependent transcription at G-less cassettes and could, thus, be extended even if Rpa12.2 is lacking. In contrast recovery from deep backtrack depths requires the cleavage supporting activity of Rpa12.2 (Lisica et al., 2016). It is possible that such deep backtrack depths are facilitated in the presence of the dimerization module. Taken together, it is likely that the dimerization module plays an important role to maintain stalled transcription complexes, which can only resume elongation if the cleavage supporting activity of Rpa12.2 is present.

### Cooperation between the heterodimer and full length Rpa12.2 prevents NTP mis-incorporation and is required to elongate stalled transcripts

To confirm that the dimerization domain influenced transcription fidelity and stalling of elongation complexes by cooperating with Rpa12.2, transcription reactions using tailed templates with G-less cassettes were performed (Fig. 5A). As stated above, transcription reactions using tailed templates yielded always more transcripts with misincorporated nucleotides (readthrough transcripts) than promoter-dependent transcription reactions. As expected, addition of Rpa49CT to ΔRpa49 Pol I, ΔRpa12 Pol I and Rpa12ΔC Pol I stimulated readthrough transcription (Fig. 5A, compare lane 2 with 1, lane 6 with 5 and lane 10 with 9). Addition of the dimerization domain to ΔRpa49 reduced slightly readthrough transcripts but did not significantly change the transcript pattern of the other two mutant polymerases (Fig. 5A, compare lane 3 with 1, lane 7 with 5 and lane 11 with 9 and profile analysis aside). Addition of the dimerization domain to a Rpa49CT-stimulated ΔRpa49 Pol I dependent reaction largely suppressed readthrough transcription, confirming that cooperation of both domains of Rpa34.5/Rpa49 are required for correct and efficient NTP incorporation (Fig. 5A, compare lane 4 and 2, and the pattern profile aside). In contrast, addition of the dimerization domain to Rpa49CT-stimulated transcription of either ΔRpa12 or Rpa12ΔC resulted in not such pronounced changes of the transcript pattern (compare lane 8 with 6 and lane 12 with 10 and pattern profile aside). This indicates that the RNA cleavage supporting activity of Rpa12.2 together with the dimerization domain or the complete heterodimer is required to prevent NTP misincorporation at the G-less cassette.

**Fig. 5.**
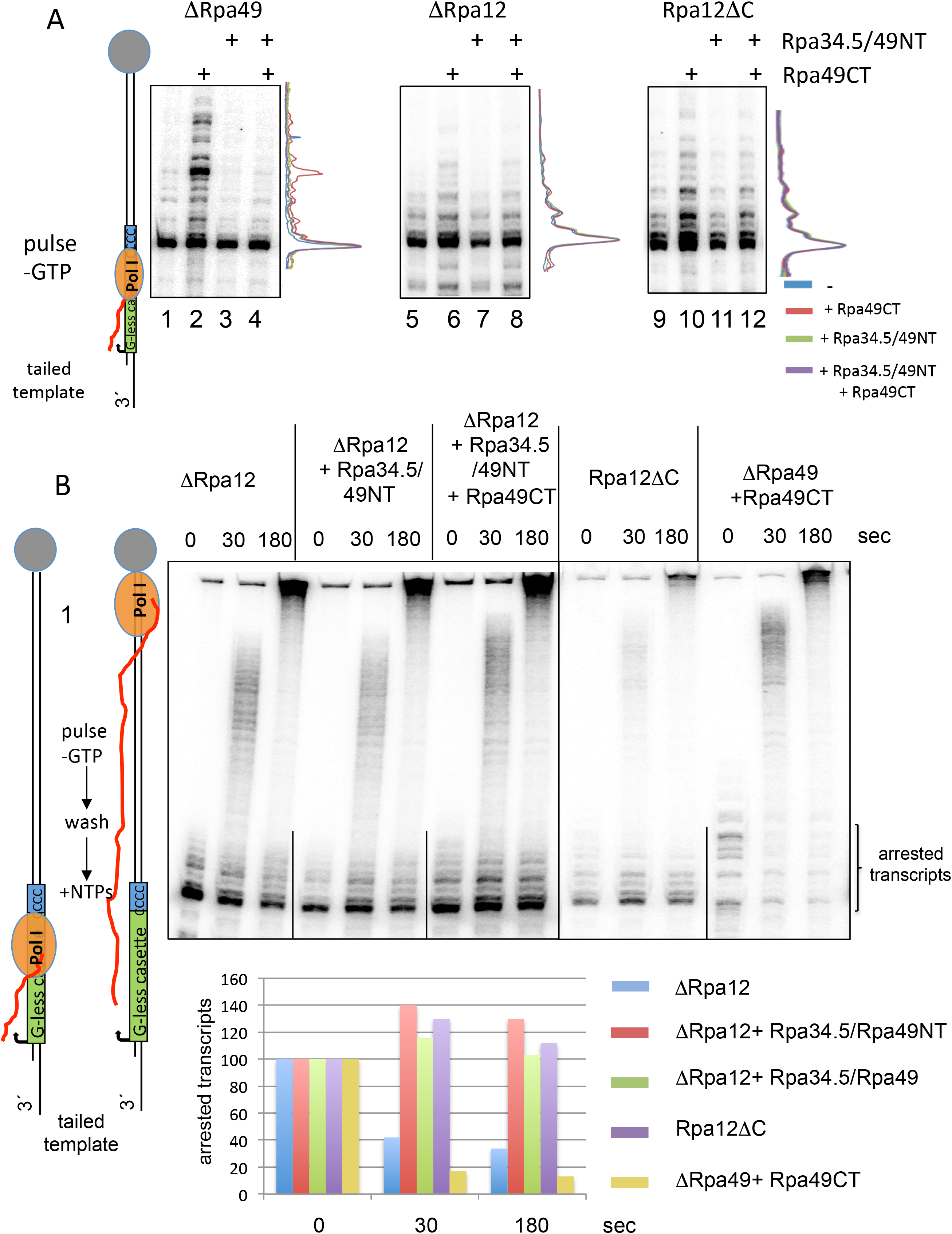
Cooperation between Rpa12 and the heterodimer Rpa34.5/Rpa49 promotes correct incorporation of nucleotides and re-extension of arrested transcripts. A) The dimerization domain supports transcription fidelity only if the RNA cleaving activity of Rpa12.2 is present. Transcription reactions were performed using 5 nM mutant enzymes in the presence or absence of Rpa34.5/Rpa49 domains as indicated (each 40nM). Pulse labelling on tailed templates containing a G-less cassettes occurred in the presence of ^32^P-CTP, ATP and UTP. Transcripts were analyzed on a 6% polyacrylamide gel containing 6M urea. Superimposed transcript profiles of the 4 single reactions are depicted near each section. The profiles were normalized to the band representing correct arrested transcripts. Note that Rpa34.5/Rpa49NT reduced mis-incorporation of nucleotides, when it was added to ΔRpa49 + Rpa49CT, which contains Rpa12.2. In contrast, reduction of NTP mis-incorporation was less effective after addition of RPA34.5/Rpa49NT to mutants lacking either Rpa12.2 or Rpa12.2CT. Experiments were conducted at least in duplicate. B) The dimerization domain without Rpa12.2 produces stalled transcripts at the G-less cassette which cannot be extended. Transcription complexes containing ^32^P-labelled RNA were generated on tailed templates using the Pol I mutants and protein fractions as indicated and stalled at the G-less cassette (10 min pulse labelling without GTP). Complexes were washed once, NTPs (200 μM end concentration) were added and the reactions were stopped after 30 and 180 seconds respectively. Transcripts were analyzed on a 6% polyacrylamide gel containing 6 M urea. (Note, the radiolabeled band on top of all lanes is due to non-specific labelling). Quantification of the arrested transcripts (in brackets) is depicted at the diagram below. (Experiment is representative for several replicates n>3, see also Suppl. Fig. 4)

Our previous analysis suggested that either WT Pol I or enzymes lacking the dimerization domain could efficiently resume elongation of arrested transcripts after the G-less cassette (see Fig. 1–4). As published for Pol I and II transcription (Izban and Luse, 1993) (Lisica et al., 2016) and described above, different backtracking depths of ternary elongation complexes can occur. Accordingly, polymerases can recover from shallow backtracks by 1D diffusion whereas deeper backtracks require RNA cleavage activity. Therefore, stalled complexes of Pol I mutants lacking the dimerization domain might remain elongation-competent no matter whether the enzyme can cleave RNA or not. In contrast, the presence of the dimerization domain could support a larger backtrack depth which then requires RNA cleavage activity to restart elongation. To find out whether the presence of the dimerization domain Rpa34.5/Rpa49NT is sufficient to prevent chain elongation if RNA cleavage cannot take place, we asked which proportion of arrested transcripts can be extended in the presence and absence of the dimerization module Rpa34.5/Rpa49NT when Rpa12.2 is missing. Pulse experiments were performed using tailed templates in the presence of ^32^P-CTP, ATP and UTP, ternary complexes were washed and after addition of nucleotides the amount of transcripts at the G-less cassette which were not extended were analysed after 30 and 180 seconds (Fig. 5B/ Suppl Fig. 4A). As already shown in Fig. 2 and 4, a significant proportion of ΔRpa12.2 Pol I was able to extend stalled transcripts. A similar effect was observed for ΔRpa49 Pol I substituted with Rpa49CT. In contrast, addition of either the complete heterodimer Rpa34.5/Rpa49 or the dimerization domain Rpa34.5/Rpa49NT revealed that a major population of the transcripts remained arrested. The same was true for the Rpa12ΔC mutant. Promoter-dependent transcription reactions using the same experimental approach showed comparable results with the exception that the influence of Rpa34.5/Rpa49NT could not be tested in promoter-dependent assays (Suppl. Fig. 4B). In summary, these experiments suggest that the dimerization domain is important to establish a transcriptional state, in which Rpa12.2 dependent cleavage is required for efficient extension of arrested transcripts. On the other hand, Pol I mutants lacking the heterodimer or only the dimerization domain generated much less non-extendable transcripts at the G-less cassette no matter whether the RNA cleaving activity of Rpa12.2 was available or not. Probably, these mutants bypass the backtracking/cleavage process of the wildtype enzyme when the enzyme is stalled during elongation.

### Pol I and Pol III which contain lobe-binding modules but not Pol II transcribe through an assembled nucleosome

In contrast to Pol I and Pol III the lb-module which is formed by subunits RPA34.5/Rpa49 and Rpc37/Rpc53, respectively, is absent in Pol II. It was previously published that purified Pol III can transcribe through a nucleosome (Studitsky et al., 1997) whereas Pol II is inhibited (Izban and Luse, 1991) without inclusion of additional factors (Bondarenko et al., 2006). Stimulation of Pol II transcription through nucleosomes was achieved in the presence of TFIIS and the elongation factor FACT (Bondarenko et al., 2006) and when TFIIS and TFIIF were combined (Luse et al., 2011). In mammalian Pol I and Pol III transcription FACT was described to facilitate chromatin transcription (Birch et al., 2009). We wanted to know, if the three purified nuclear core RNA polymerases from yeast without additional factors differ in their ability to transcribe nucleosomal templates. A single nucleosome was assembled by salt dialysis on a tailed template containing a 601 nucleosome positioning sequence (see Suppl. Fig. 5 and Material and Methods). This set-up allowed us to analyse the efficiency of affinity purified Pol I to synthesize RNA from nucleosomal templates in comparison to the same nucleosome-free templates without inclusion of initiation factors. To quantify the efficiency of RNA synthesis through an assembled nucleosome, transcription reactions were always performed in the presence of a nucleosome-free either shorter or longer reference template (Fig. 6 and Fig. 7). Reactions were stopped after 30 min. Pol I, Pol II and Pol III were purified according to a previously published protocol which allowed the identical one-step affinity purification procedure using the same buffer conditions. The purified enzymes were free of detectable cross-contamination of transcription factors or RNA synthesizing enzymes (Merkl et al., 2014). Transcription reactions were performed each with 7.5 nM of the respective polymerases and identical buffer conditions (50 mM KoAc), which had been tested to allow transcription by all three polymerases. All three polymerases were able to transcribe reference templates and the 601 sequence containing templates without nucleosomes quite efficiently (Fig. 6, lanes 2, 4, 5, 7, 9, 10, 12, 14, 15). Differences in signal intensities between the generated RNAs were due to the buffer conditions which were not optimal for all enzymes. The presence of a nucleosome on the DNA template reduced the efficiency of transcription. However, the level of inhibition differed significantly between the three polymerases. Whereas a major fraction of Pol I and Pol III could transcribe the nucleosomal 601 sequence (Fig. 6A, lanes 1, 3, 11, 13), full length transcripts from nucleosomal templates were almost not detectable in Pol II-dependent reactions (lanes 6 and 8). Upon longer exposure times an additional transcript became visible in lanes 6 and 8 which length corresponded to the size from the start site to the 5’end of the positioned nucleosome (Suppl Fig. 6). Apparently, abortive or arrested transcripts were generated when the enzymes encountered a nucleosome. This indicated that at least a minor fraction of Pol II initiated transcription from the tailed template but was arrested or displaced upon collision with a nucleosome. Quantification revealed about 40% reduction of RNA synthesis by Pol I and Pol III in the presence of a nucleosome and almost complete loss of Pol II-derived transcripts (Fig. 6). These results suggest that Pol I and Pol III which both contain additional heterodimeric subcomplexes (Rpa34.5/Rpa49 and Rpc37/Rpc53, respectively) in comparison to Pol II, have intrinsic activities to transcribe through a nucleosomal barrier.

**Fig 6.**
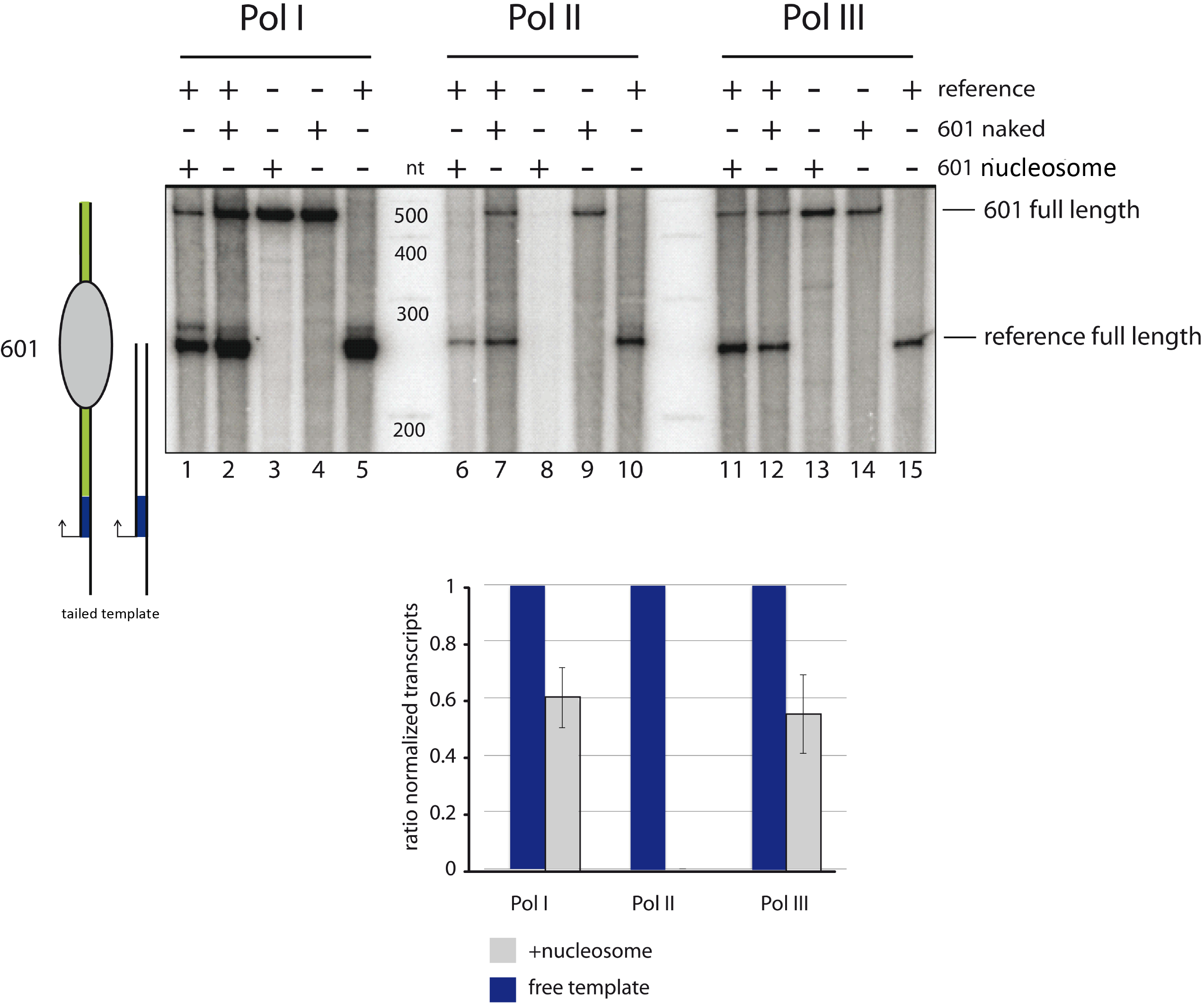
Purified Pol I and Pol III transcribe through a nucleosome on tailed templates more efficiently than Pol II. Nucleosomes were assembled on tailed templates containing a 601 nucleosome positioning sequence (601) (derived from plasmid 1253) as described in material and methods (see also Suppl Fig. 5). Transcription reactions were performed in the presence of a nucleosome-containing and a nucleosome-free reference template. The cartoon on the left indicates the length of the transcripts derived from the nucleosomal and reference template. *In vitro* transcription assays were performed as described using 7.5 nM of purified RNA polymerases, identical buffer conditions and 10 nM of template 601 with or without nucleosomes and the reference template containing neither nucleosomes nor the 601 sequence. The cartoon on the left indicates the length and the anticipated position of the nucleosome of the transcribed templates. *Lower panel* Quantification of the experiment. Signal intensities of 601 full length transcripts (lanes 1/2, 6/7 and 11/12) were normalized to reference full length transcript signal intensities in the respective lanes. Then, the ratio of the normalized 601 full length transcript signal intensities in presence and absence of the nucleosome was calculated. Experiments were conducted at least in duplicate and mean values were plotted including error bars of the standard deviation. Quantification of the putative Pol II transcript after 24h exposition did not result in a signal above background level. Length of reference transcript 240 nt Length of the 601 sequence 468 nt

**Fig. 7.**
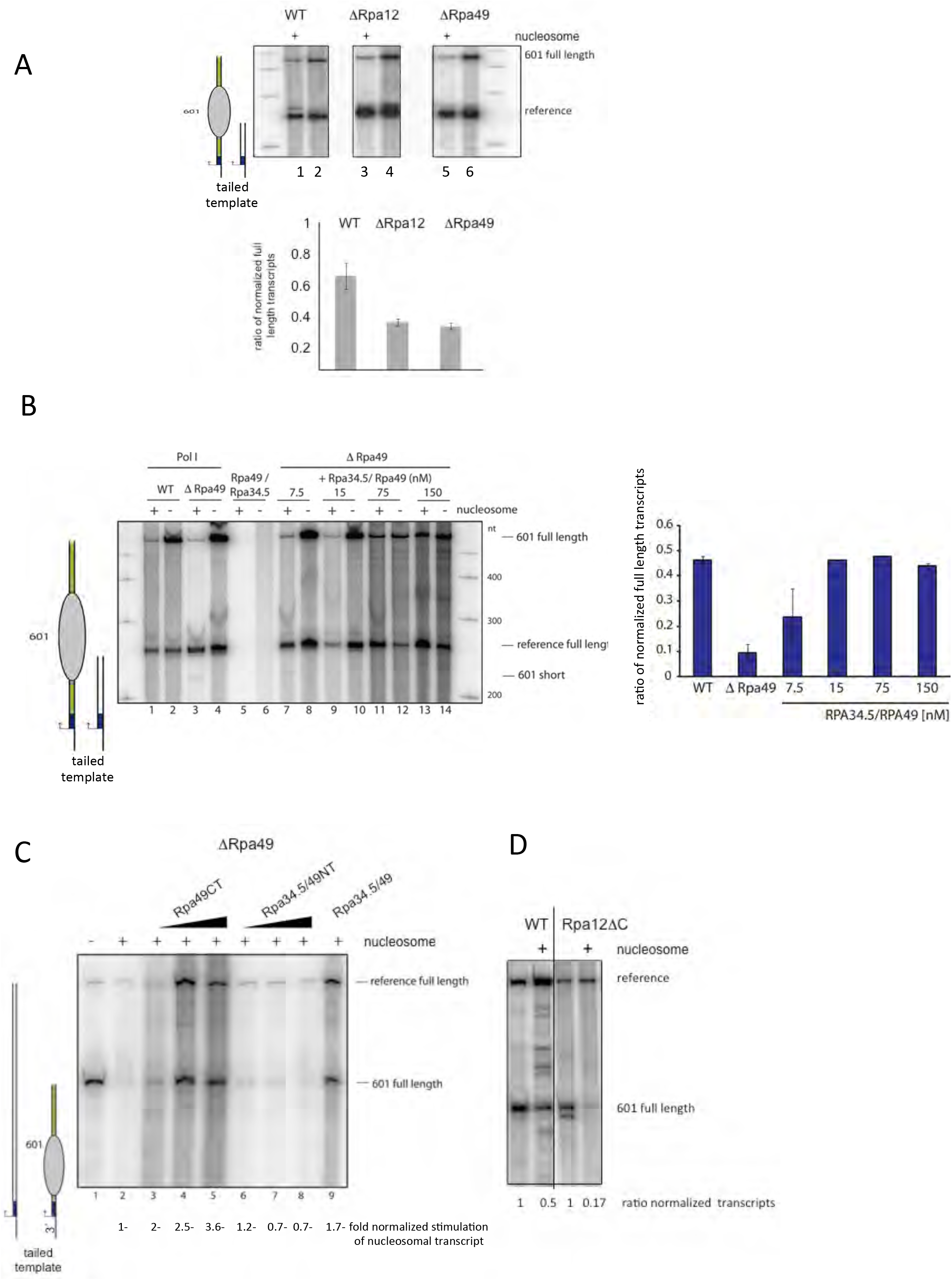
Efficient transcription through nucleosomes requires Rpa34.5/Rpa49 and Rpa12.2. A) Transcription reactions were performed on tailed templates using 20 nM ΔRpa49 or ΔRpa12 as in Fig. 6. The cartoon on the left indicates the length of the transcripts derived from the nucleosomal and reference template. Transcripts were analyzed on a 6% polyacrylamide gel containing 6 M urea. Experiments were conducted in duplicate. B) The heterodimeric subunit Rpa34.5/Rpa49 of Pol I is required for efficient transcription through a nucleosome in tailed template assays. *Upper panel.* Transcription reactions were performed using 15 nM Pol I lacking subunits Rpa49 and Rpa34.5 (ΔRpa49), lanes 1 and 2). Increasing amounts of purified recombinant Rpa34.5/Rpa49 heterodimer were added to ΔRpa49-Pol I in reactions analysed in lanes 5 to 12. Reactions analysed in lanes 3 and 4 were performed using 150 nM Rpa34.5/Rpa49 heterodimer, but no polymerase. (601 short indicates the position of a stalled transcript at the 5’end of a nucleosome; see also Suppl Fig. 4). *Right panel.* Quantification from two independent experiments using the same preparation of ΔRpa49. C) Effects of Rpa49 domains on transcription of a nucleosomal tailed template. Add-back experiments to ΔRpa49 were performed using 25, 50 and 75 nM of either purified recombinant heterodimeric subunits Rpa49CT, Rpa34.5/Rpa49NT, or 50 nM Rpa34.5/Rpa49. Concentration of ΔRpa49 Pol I was 10 nM. In contrast to the previous tailed template transcriptions the length of the templates were switched. Template concentration was 7.5 nM. The transcripts from the 601 and reference template were 240 nt and 468 nt, respectively. This template combination gave less non-specific transcripts in the range between 250 nt and 500 nt in the presence of Rpa49CT. The stimulation of transcription through nucleosomes in reference to the stimulation of transcription on the nucleosome-free DNA is shown underneath. The same preparation of WT Pol I and ΔRpa49 was used. D) Efficient transcription through nucleosomes requires the C-terminal domain of Rpa12.2. Tailed template transcription was performed as in C, using 10 nM WT Pol I or Rpa12.2AC. Quantification was performed as in Fig. 6.

### Pol I transcription of nucleosomal templates *in vitro* is compromised in the absence of the Pol I-specific heterodimer Rpa34.5/Rpa49 and when the C-terminal part of Rpa12.2 is missing

In Pol II-dependent transcription TFIIF together with the cleavage stimulating factor TFIIS promotes elongation (Schweikhard et al., 2014) and RNA synthesis through nucleosomes (Luse et al., 2011). Since Rpa34.5/49 and Rpa12.2 share structural features with TFIIF and TFIIS, one could hypothesize that these subunits might facilitate transcription through a nucleosome. Therefore, we investigated whether absence of Rpa12.2 with and without the Rpa34.5/49 heterodimer had an impact on transcription of a nucleosomal template. Consistent with a role of these subunits in transcription through a nucleosome, both, ΔRpa12.2 Pol I and ΔRpa49 Pol I could not as efficiently transcribe nucleosomal templates as the WT enzyme (Fig. 7A, lanes 3, 4 and 5, 6). Addition of increasing amounts of the purified heterodimer restored the ability of ΔRpa49 Pol I to transcribe through a nucleosome (Fig. 7B, lanes 7–14, and quantification), while the purified heterodimer alone contained no RNA synthesizing activities (lanes 5 and 6). To determine the domains of the Rpa34.5/Rpa49 heterodimer which are required to stimulate transcription through a nucleosome, either purified dimerization domain Rpa34.5/49NT or the tandem winged helix-containing Rpa49CT were added to a ΔRpa49 Pol I-dependent transcription reaction. Addition of the complete heterodimer or Rpa49CT alone stimulated both, transcription of the reference template, and transcription of the nucleosomal template. (The level of stimulation of the reference template varied in dependency on the quality of the preparation of the nucleosomal DNA matrix). The best recovery of transcriptional activity on the nucleosomal template in reference to the nucleosome-free template was obtained adding back either the entire heterodimer Rpa34.5/49 or Rpa49CT to the mutant enzyme (Fig. 7C, lanes 3–5 and 9). Adding the dimerization domain Rpa34.5/Rpa49NT alone to ΔRpa49 Pol I did not show significant enhancement of transcription (Fig. 7C, left panel, compare lanes 3–5 and 6–8). RNA synthesis through nucleosomal templates using promoter-dependent transcription assays showed comparable results (Table 1).

**Table 1.** Addition of the complete heterodimer RPA34.5/49 or Rpa49CT to ΔRpa49 results in stimulation of promoter-dependent transcription on nucleosomal templates. Signal intensities of 601 full length transcripts were normalized to reference full length transcript signal intensities in the respective lanes. Then, the ratio of the normalized 601 full length transcript signal intensities in presence and absence of the nucleosome was calculated. The value for the ΔRpa49 enzyme was set to 1 and the stimulation adding the different Rpa34.5/49 domains (150nM) was calculated. Mean values of each two experiments are shown.

|  |  |
| --- | --- |
| ΔRpa49 | 1 |
| + Rpa34.5/49 | 2.7 +/- 0.6 |
| + Rpa34.5/49 NT | 1.2 +/- 0.1 |
| + Rpa49 CT | 2.0 +/- 0.3 |

To find out whether the RNA cleavage-stimulating activity of Rpa12.2 contributes to transcription through nucleosomes, transcription assays on tailed templates were performed using WT Pol I and Rpa12ΔCT. In the presence of nucleosomes transcription using Rpa12ΔCT was much stronger inhibited than with WT Pol I, suggesting that - in addition to the heterodimer - RNA cleavage activity is also required for efficient passage through a nucleosome. In summary, the dimerization module which is a constituent of the Pol I lb-module, the C-terminal part of Rpa49 and the C-terminal part of Rpa12.2 seem to cooperate to support Pol I transcription of both, nucleosomal and nucleosome-free templates.

## Discussion

### Comparison of Rpa34.5/49 and Rpa12.2 with their Pol II counterparts

The functional properties of the heterodimeric Rpa34.5/Rpa49 complex and Rpa12.2 show significant similarities to the transcription factors TFIIE, TFIIF, and TFIIS, their counterparts in Pol II dependent transcription (Kuhn et al., 2007, Geiger et al., 2010). TFIIE and TFIIF have specific roles in initiation and elongation and are not components of the core enzyme. Similar to Rpa49CT and, Rpa34.5/Rpa49NT, TFIIE and the dimerization module of TFIIF bind each to opposite sides of Pol II, respectively, and flank the promoter DNA. Both Pol II factors are parts of an intricate transcription initiation network containing TFIIB, TBP and domains of the Pol II core enzyme which are thought to encircle, retain and open the DNA (He et al., 2016, Plaschka et al., 2016). The Tfa2 tWH domain of TFIIE corresponds to Rpa49Ct, but is only important for transcription initiation. Tfa2 spans the polymerase cleft and is positioned close to the site of DNA melting (Grunberg et al., 2012). Recent structural data revealed a similar position for Rpa49CT which stabilizes the upstream DNA during transcription (Pilsl et al., 2016a, Tafur et al., 2016, Jennebach et al., 2012). The Rpa49 linker, which is part of Rpa49CT, connects the tWH domain of Rpa49 with the dimerization module, thereby spanning the active site cleft and contacting the coiled coil (Tafur et al., 2016, Han et al., 2017). Based on the structures of different Pol I fractions, it was suggested that closing of the Pol I clamp and establishing extensive contacts with the RNA and the upstream DNA correlate with the stabilisation of the Rpa49CT linker region over the cleft. In correlation to TFIIE function in initiation, this might cause the stabilisation of the tWH domain of Rpa49CT during transcription initiation (Tafur et al., 2016, Han et al., 2017, Engel et al., 2013). Contrary to TFIIE, Rpa49CT accompanies the elongating enzyme (Beckouet et al., 2008). Therefore, clamp closing through Rpa49CT could also establish an elongation promoting Pol I complex which is dedicated for the fast incorporation of nucleotides.

TFIIF the counterpart of Rpa34.5/49NT was described as initiation and elongation factor. This is in contrast to Rpa34.5/49NT which is neither required to stimulate initiation *in vitro* (Pilsl et al., 2016a) nor for Pol I loading onto the promoter (Beckouet et al., 2008). TFIIF predominantly associates with Pol II in promoter-proximal regions (Mayer et al., 2010) (Krogan et al., 2002). In addition, it accompanies at least transiently Pol II during RNA chain elongation (Rani et al., 2004, Mayer et al., 2010, Krogan et al., 2002, Cojocaru et al., 2008). It was shown, that TFIIF is involved in very early stages of elongation (Yan et al., 1999, Cheng and Price, 2007, Ujvari et al., 2011) and that it increases transcription rates *in vitro* (Renner et al., 2001, Izban and Luse, 1992, Zhang et al., 2003), at least in part by suppressing pauses. Presumably, TFIIF promotes transcription elongation together with the RNA cleavage stimulating factor TFIIS in a synergistic manner (Schweikhard et al., 2014, Zhang et al., 2003, Zhang and Burton, 2004, Zhang et al., 2005), and the impact of both proteins is important to increase transcription through nucleosomes *in vitro* (Luse et al., 2011). These elongation related features resemble those of the Pol I dimerization module and its interaction with the RNA cleavage stimulating factor Rpa12.2.

### Functions of the heterodimeric Rpa34.5/Rpa49 complex and interplay with Rpa12.2

The dimerization module Rpa34.5/49NT and the N-terminal part of Rpa12.2 form the polymerase lobe-binding module which is not present in the Pol II enzyme. Our studies underline that these subunits functionally cooperate. Efficient RNA chain elongation was dependent on the Rpa34.5/Rpa49 complex. Apparently, this complex is important for optimized Pol I processivity, elongation speed, pausing and RNA cleavage events. All elongation features depended on the cooperation of both domains. *In vitro* transcription indicated that Rpa49CT increased elongation speed, but lowered significantly Pol I processivity and transcription fidelity, if the dimerization module Rpa34.5/Rpa49NT was lacking. This suggested that the tandem winged helix (tWH) of Rpa49CT promotes the core enzyme to incorporate nucleotides at higher rate, but at the cost of less accuracy. Interaction of the tWH with the dimerization module reduced the speed of RNA-synthesis and misincorporation of nucleotides, and, importantly, increased Pol I processivity.

In the absence of the dimerization module, the presence of Rpa12.2 had neither influence on speed, transcription fidelity, extension of arrested transcripts and processivity (this manuscript), nor on RNA cleavage activity (Kuhn et al., 2007, Geiger et al., 2010). This underlined that Rpa12.2 function depended on the neighbouring dimerization domain and confirmed previous suggestions that the cleavage stimulatory effect of the heterodimer is indirect and due to its proximity to Rpa12.2 (Jennebach et al., 2012). Taken together, these findings suggest that the dimerization domain containing lb-module is required to integrate Rpa12.2- and Rpa49CT-dependent activities into the transcribing enzyme which finally ensures highly processive and accurate transcription. It could be that a similar interplay is responsible for the reported synergism between TFIIF with TFIIS which ensures Pol II proofreading (Thomas et al., 1998, Jeon and Agarwal, 1996, Schweikhard et al., 2014).

### The interplay of the lobe binding subunits with stalled transcription complexes

If RNA polymerases encounter a barrier, the enzyme stalls, backtracks and arrests (Luse et al., 2011, Lisica et al., 2016, Cheung and Cramer, 2011). Rapid relief of arrest requires RNA cleavage stimulating proteins like TFIIS, Rpa12.2 or the Pol III subunit C11 and cleavage of the 3’ end of the RNA which is disposed from the active center. In the absence of RNA cleaving activity, NTPs were misincorporated in Pol I-dependent transcription. If either the dimerization module or the C-terminus of Rpa12.2 was lacking, transcription did not stop precisely at the G-less cassette. This resembles recent findings that Rpa12ΔC Pol I in contrast to WT Pol I can slowly incorporate nucleotides opposite to a cyclobutene pyrimidine dimer (Sanz-Murillo et al., 2018). However, although RNA cleavage activity is largely reduced in the absence of the dimerization domain (ΔRpa49, ΔRpa12.2)(Kuhn et al., 2007), a significant fraction of these enzymes can resume RNA chain elongation after ternary complexes were paused at G-less cassettes (Fig. 1–4). It is possible that extension of RNA occurs if *i*) the enzyme randomly diffuses forward to realign the RNA 3’end or *ii*) complexes do not enter the deep backtracked state which requires RNA cleavage for elongation when they encounter the G-less cassette arrest or other obstacles as it was previously reported for Pol II transcription *in vitro* (Chang and Luse, 1997). The presence of Rpa49CT could enforce such a non-arrested stage. Interestingly, Pol I transcription is not efficiently terminated in ΔRpa*12* strains (Prescott et al., 2004), which could also be due to non-arrested complexes at the termination protein Nsi1 (Reiter et al., 2012). The dimerization module resides on the polymerase lobe in close vicinity to subunit Rpa12.2, which is required for RNA cleavage of backtracked Pol I. If lack of the dimerization module destabilizes Rpa12.2 such that it cannot stably access the active center, it is conceivable that such an enzyme *i*) is not able to support cleavage reactions and *ii*) may not enter a locked state with a 3’ RNA displaced from the active center. One consequence would be a reduced transcription fidelity. On the other hand, it is possible that the dimerization domain supports the generation of deep backtracked intermediates in specific situations which can only resume elongation if RNA cleavage can take place. It will be interesting to see if incorporation of a wrong nucleotide can facilitate such a Rpa34.5/Rpa49NT driven backtrack. Very recently a manuscript was submitted in which it was suggested that the association of the Rpa34.5/Rpa49 is regulated by the C-terminal domain of Rpa12.2 (Tafur et al., 2018). If (transient) dissociation of the dimerization domain by Rpa12.2 is important to restart RNA synthesis, it is conceivable that a Pol I mutant lacking the Rpa12.2 C-terminus but including the dimerization domain has problems to enter the elongation mode.

### How to transcribe through a nucleosome?

Transcription through nucleosomes is a special challenge for eukaryotic RNA polymerases since intrinsic stalling and backtracking is more pronounced in front of a physical barrier like a nucleosome. Furthermore, the nucleosomal DNA recoils on the octamer, locking the enzyme in the arrested state (Gaykalova et al., 2015). Rapid relief of arrest requires transcription factor TFIIS and cleavage of the 3’ end of the RNA which is disposed from the active center of Pol II. TFIIF was reported to support TFIIS-mediated *in vitro* transcription through nucleosomes (Luse et al., 2011). Both factors were not associated to Pol II, when Pol II was affinity-purified using our stringent conditions. Thus, Pol II was not able to transcribe through nucleosomes. In contrast Pol I and Pol III which have the counterparts of TFIIF and TFIIS, the subunits Rpa34.5/Rpa49, Rpa12.2, and Rpc37/Rpc53, Rpc11, respectively, tightly associated, were able to pass through nucleosomes. For Pol I, efficient nucleosomal transcription depended on the presence of the heterodimeric Rpa34.5/Rpa49 complex. Apparently, both modules Rpa34.5/Rpa49NT and Rpa49CT as well as the C-terminal part of Rpa12.2 contributed to the ability to transcribe through nucleosomes, suggesting that both cleavage activity and Pol I binding to DNA (mediated by Rpa49CT) support passage through nucleosomes. In the absence of the dimerization module, addition of Rpa49CT alone improved Pol I movement through nucleosomes. As mentioned above, it is possible that such an enzyme may not enter a locked state with a 3’ RNA displaced from the active center, suggesting that elevated speed generated by Rpa49CT compensate for the missing backtracking and RNA cleavage cycle. Whether or not the nucleosome is evicted by Pol I or the DNA is transiently uncoiled during elongation remains to be determined. So far, the sensitivity of our assays applied was not sufficient to allow clear-cut conclusions.

For passage through nucleosomes uncoiling of the template from the histone octamer while maintaining the polymerase in the transcriptionally competent state are the most important features. Histone chaperone FACT or elongation factors Spt4/5 or Paf1c are examples of Pol II factors that facilitate DNA unwrapping from H2A/H2B dimers to partially disassemble nucleosomes (Belotserkovskaya et al., 2003, Hsieh et al., 2013, Crickard et al., 2017, Kim et al., 2010). The same factors were also reported either to support Pol I elongation (Zhang et al., 2009, Zhang et al., 2010, Schneider et al., 2006) or passage through nucleosomes (Birch et al., 2009). None of these factors was present in the tailed template assays performed in this study. But, it is possible that a combinatory effort between Pol I subunits and elongation factors further support transcription through nucleosomes.

### *In vitro* and *in vivo* phenotypes

In contrast to Pol II genes, Pol I transcribed genes are heavily packed with elongating polymerases which do not leave much space for assembled nucleosomes (Merz et al., 2008, Conconi et al., 1989). It was previously shown that Pol I transcription is required to achieve an open chromatin state in which the amount of DNA-bound nucleosomes is strongly reduced (Merz et al., 2008, Wittner et al., 2011). According to our *in vitro* studies the Pol I heterodimer Rpa34.5/Rpa49 and the Rpa12.2 C-terminus are involved in this process. On the other hand mutant strains which are deleted in the heterodimer or RPA12.2 are viable (Liljelund et al., 1992, Nogi et al., 1993) (Gadal et al., 1997). This suggests that these mutant polymerases can still open the closed chromatin state (Albert et al., 2011) and that other factors of the Pol I transcription machinery contribute to the opening process. Such candidates could be the above mentioned FACT (Birch et al., 2009) or Paf1c (Zhang et al., 2010). Future structural and functional investigations are necessary to elucidate mechanisms how stalled Pol I complexes can bypass transcription obstacles *in vivo* with and without removal of a displaced RNA 3’end. Taken together our studies underline that the specific subunit composition of Pol I adds to the functional specialisation of the enzyme to transcribe the rRNA gene as efficiently as possible.

## Materials and Methods

### Yeast strains, plasmids, oligonucleotides and construction of transcription templates

Oligonucleotides, plasmids and yeast strains used in this work are listed in suppl. Tables 1 2 and 3. Molecular biological methods and transformation of yeast cells were performed according to standard protocols (Sambrook et al., 1989, Burke et al., 2000, Schiestl and Gietz, 1989). The generation of transcription templates is described in the supporting information. Plasmid sequences are available upon request. For the experiments in this study, yeast cells were grown at the indicated temperatures in either YPD (2% w/v) peptone, 1% (w/v) yeast extract, 2% glucose).

### Purification of proteins

Wild-type RNA polymerases I, II and III (Pol I, II and III) were purified from yeast strains y2423 (Pol I), y2424 (Pol II), y2425 Pol III, (suppl. Table 1) via the protein A affinity tag according to (Merkl et al., 2014). Purification of mutant Pol I y2670 (ΔRpa49), y2679 (ΔRpa12) and y2679 transformed with plasmid 1839 (Rpa12ΔCT) was according to (Pilsl et al., 2016a). Recombinant Rpa34/Rpa49, the dimerization domain Rpa34.5/Rpa49NT (Rpa34/(Rpa49 aa 1–110) and Rpa49CT (aa 110–426) were purified from *E.coli* according the protocol of (Pilsl et al., 2016a). The absolute efficiency of single protein fractions in transcriptional activity varied slightly from preparation to preparation. Peculiar attention concerned the expression and purification of ΔRpa49 Pol I (y2670). To avoid the observed generation of suppressors this strain was first cultivated in galactose, which allowed the expression of Rpa49 and then shifted to glucose for 24 hours to shut off Rpa49 expression. Although depletion of subunits A49 and A34.5 was monitored by Coomassie-stained SDS PAGE and western blotting the transcriptional activity varied slightly in different preparations. This could be due to different levels of remaining Rpa49 or upcoming suppressors. Therefore, statistical evaluation using biological replicates (which means different protein preparations from independently cultivated strains) was not applicable. Recombinant Rpa34/Rpa49 and its purified subdomains tend to loose activity when they are stored. Reliable results were obtained using freshly prepared protein fractions.

### Transcription elongation assays

Promoter-dependent elongation assay. Pulse-labelling of G-less transcripts on promoter-containing templates and subsequent analysis were performed according to (Tschochner, 1996, Pilsl et al., 2016a) with some modifications. If elongation should be measured during 5 time points including the starting point at time point 0, the following assay was performed. 1.5 ml reaction tubes (Sarstedt safety seal) were placed on ice. 5 × 0.5 to 1 μl template (50 - to 100 ng DNA) (templates 2148 or 2313) immobilized to magnetic beads was added, which corresponds to a final concentration of 5 - 10 nM per transcription reaction (25 μl reaction volume). 5 × 1-2 μl CF (0.5 to 1 pmol/μl; final concentration 20 - 40 nM) and 5 × 1-3 μl Pol I-Rrn3 (final concentration 4 - 12 nM) were added to the tube. Where indicated, domains of the Rpa34.5/49 heterodimer were added before transcription was started (final concentration 50 nM). 20 mM HEPES/KOH pH 7.8 were added to a final volume of 5 × 12.5 μl. Transcription was started adding 5 × 12.5 μl transcription buffer 2 x without GTP (40 mM HEPES/KOH pH 7.8, 20 mM MgCl_2_, 10 mM EGTA, 5 mMDTT, 300 mM potassium acetate, 0.4 mM ATP,, 0.4 mM UTP, 0.02 mM CTP, 0.3 μCi ^32^P-CTP. The samples were incubated at 24°C between 5 and 30 min at 400 rpm in a thermomixer. After placement on ice, the supernatant was separated from the magnetic beads and 2–3 washing steps using washbuffer W1 (20 mM HEPES/KOH pH 7.8, 10 mM MgCl_2_, 5 mM EGTA, 2.5 mMDTT, 150 mM potassium acetate, 0.02 mM ATP, 0.02 mM UTP, 0.02 mM CTP followed). The magnetic beads were resuspended in 150μl washbuffer W1. 25 μl were removed and added to 200 μl Proteinase K buffer (0.5 mg/ml Proteinase K in 0.3 M NaCl, 10 mM Tris/HCl pH 7.5, 5 mM EDTA, 0.6% SDS) (time point 0). To the remaining 125 μl either 12.5 μl or 1.25μ1 10xNTPs (ATP, GTP, UTP, CTP concentration each 2 mM) was added and immediately mixed. 25 or 27 μl aliquots were removed at the time points indicated and added to 200 μl Proteinase K buffer. The samples were incubated at 30°C for 15 min at 400 rpm in a thermomixer. 700 μl Ethanol p.a. were added and mixed. Nucleic acids were precipitated at −20°C over night or for 30 min at −80°C. The samples were centrifuged for 10 min at 12.000g and the supernatant was removed. The precipitate was washed with 0.15 ml 70% ethanol. After centrifugation, the supernatant was removed and the pellets were dried at 95°C for 2 minutes. RNA in the pellet was dissolved in 12 μl 80% formamide, 0.1 TBE, 0,02% bromophenol blue and 0.02% xylene cyanol. Samples were heated for 2 min under vigorous shaking at 95°C and briefly centrifuged. After loading on a 6% polyacrylamide gel containing 7 M urea and 1 x TBE RNAs were separated applying 25 watts for 30 - 40 min. The gel was dried after 10 min rinsing in water for 30 min at 80°C using a vacuum dryer. Radiolabelled transcripts are visualised using a PhosphoImager. For quantification, signal intensities were calculated using Multi Gauge (Fuji).

Tailed template elongation assay. The transcription were performed as described as above on PCR-derived templates containing biotinylated oligo downstream of the promoter in reverse direction and an oligo containing a Nb.BsmI (NEB) nicking site which allows non-specific initiation of all RNA polymerases without the addition of initiation factors.

### Transcription through nucleosomes

#### Preparation of nucleosomal templates

Chromatin was reconstituted on templates for *in vitro* transcription containing one or multiple 601 nucleosome positioning sequences (Lowary & Widom 1998). Dialysis chambers were prepared from siliconized microreaction tubes (Eppendorf) by clipping off the conical part and perforation of the cap with a red-hot metal rod (0 0.5 cm). The dialysis membrane (molecular weight cutoff 6.8kDa) was pre-wet in high salt buffer and fixed between tube and cap. The dialysis chambers were put in a floater in a bucket containing 300 ml high salt buffer, air bubbles were removed and the mixture of histones and DNA was applied to the chambers. Dialysis from 2 M NaCl to 0.23 M NaCl was performed over night at RT with constant stirring and at a low salt buffer flow rate of 200 ml/h (3 l total). Upon completion, the assembly solution was transferred to a siliconized tube, its volume was measured and chromatin was stored at 4°C. To determine the assembly success, 5 μl of the reaction were supplemented with 1μl loading buffer and analyzed on a 6% polyacrylamide, 0.4xTBE gel (Suppl Fig. 3). Gels were stained in 0.4x TBE containing ethidium bromide for 15min, washed in 0.4x TBE for 15 min and analysed on a FLA3000 imager (Fuji).

To determine optimal assembly conditions, histones were titrated to the DNA:

| **ratio**<br>**histones : DNA** | **0.4 : 1** | **0.6 : 1** | **0.8 : 1** | **1 : 1** |
| --- | --- | --- | --- | --- |
| high salt buffer | 44μl | 42.75μl | 41.5μl | 40.25μl |
| BSA (10mg/ml) | 1μl | 1μl | 1μl | 1μl |
| DNA template<br>(2μg/μl) | 2.5μl | 2.5μl | 2.5μl | 2.5μl |
| chicken histones<br>(0.8mg/ml) | 2.5μl | 3.75μl | 5μl | 6.25μl |
| total volume | 50µl | 50µl | 50µl | 50µl |

In most reaction series the ratios 0.6:1 or 0.8: 1 were used. Transcription reactions were performed as described (Pilsl et al., 2016a). Each reaction contained 50 - 100 ng of a nucleosome-containing (2316) and a 50 - 100 ng naked template (1573). Templates of approppriate sizes were generated by PCR reactions of different sizes using oligo 2115 and oligos 4231, 4232, 4233 or 4234.

## Acknowledgements

We are grateful to members of the chair Biochemistry III for critical discussion and to Elisabeth Silberhorn for technical assistance. This work was supported through grants of the Deutsche Forschungsgemeinschaft (SFB 960). There are no competing interests.

---

**Suppl Fig. 1.**
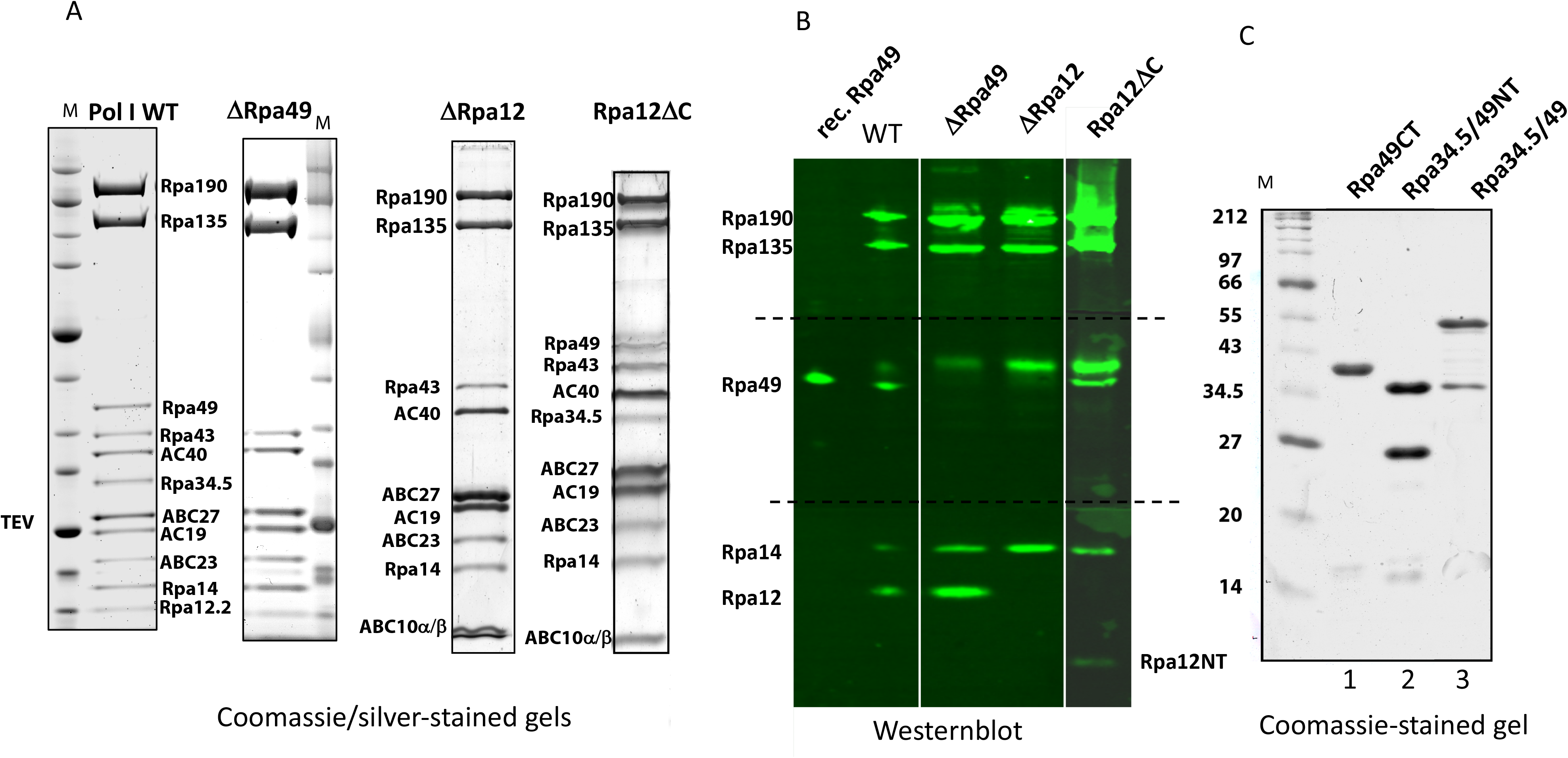
Separation of affinity-purified, ΔRpa49 Pol I ΔRpa12.2 Pol I, Rpa12ΔC and Rpa34.5/49 domain-containing fractions on SDS PAGE. A) 6 μg purified WT Pol I, ΔRpa49, ΔRpa12.2 and RPA12ΔC were separated by SDS PAGE using a 4–12% gradient gel which was stained with SimplyBlue SafeStain (ThermoFisher) or with Silver. ΔRpa49 Pol I was affinity-purified from strain yJPF162- 1a (2670) using a ProtA-TEV-tag fused to subunit A135 after 24 h depletion of subunit A49 in glucose containing medium (see Materials and Methods). Molecular weight standards (M) and Pol I subunits are indicated. B) The depletion of subunits Rpa49 and Rpa12.2 was confirmed in the affinity purified enzyme fraction by Western blotting using antibodies against Rpa49, Rpa12.2, which also recognizes Rpa14, and the two largest subunits. After blotting the membrane was cut into three pieces and equally treated with the different antibodies. Flag-tagged Rpa49 was separated as reference. (Note that Rpa49 is neither detected in ΔRpa49 nor in ΔRpa12.2 the band above Rpa49 is due to nonspecific IgG interaction). Subunit RPA12.2 is only completely depleted in strain ΔRpa12.2 (Y2679) whereas the associated N-terminal part of Rpa12.2 is detectable in the Rpa12ΔC preparation. C) Separation of recombinant A49/A34.5 domains on SDS PAGE. comparable protein amounts were separated on a 15 % SDS gel and Coomassie stained. Lane 1: Rpa49 (amino acids 110–415), 2: Rpa34.5/49 (1–186), 3: A49/A34,5 full length; Expression and purification strategy see Material and Methods and (Pilsl et al., 2016a). Fractions were used for *in vitro* transcription.

**Suppl. Fig. 2.**
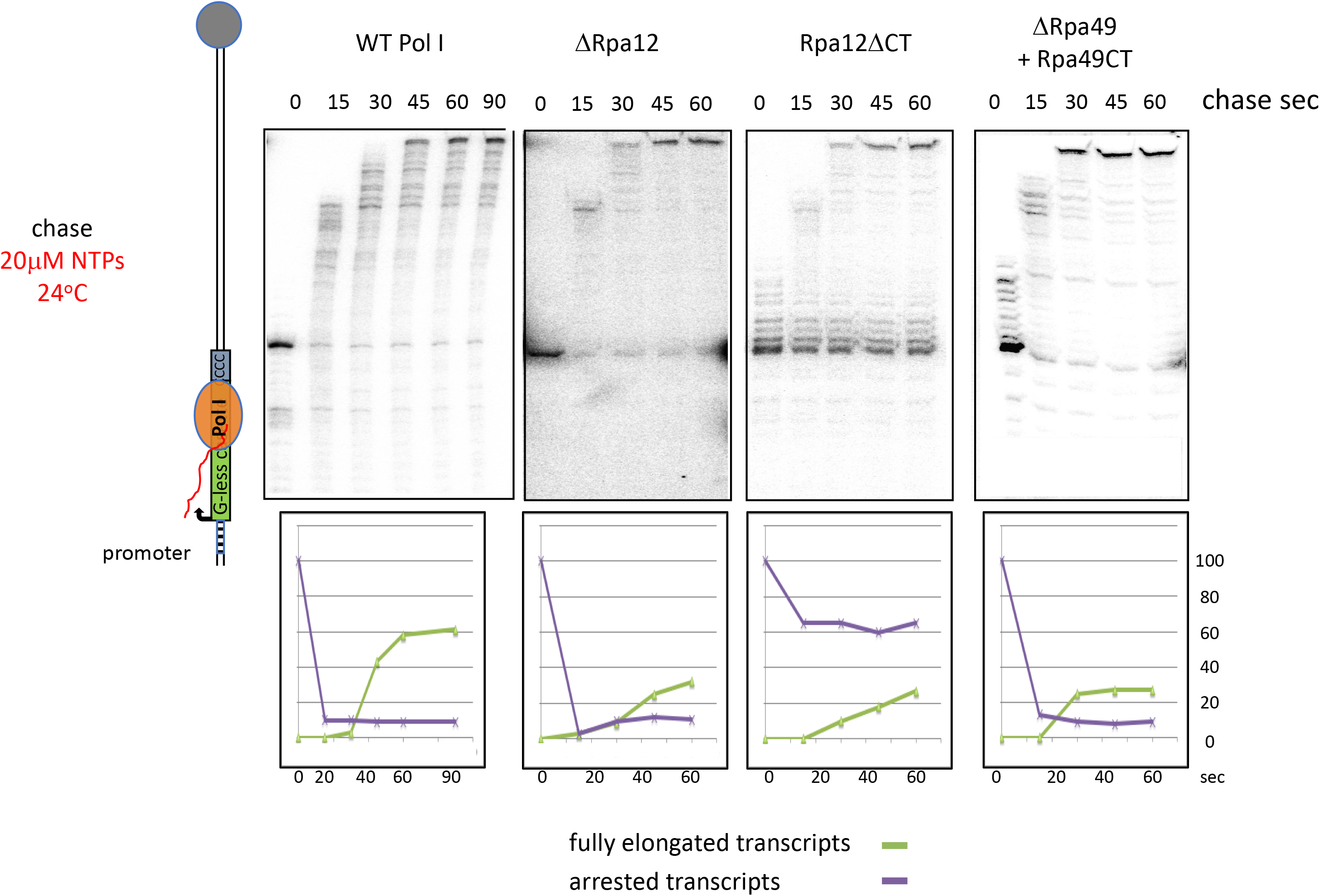
Reduction of Pol I processivity in different mutant enzymes when elongation at the G-less cassette occurs with 20 μ.M NTPs. Promoter-dependent elongation assays were performed as described in Fig. 2 with the exception that stalled transcripts were chased using 20 0M NTPs. Assays were performed using 4 nM Pol I, Rpa12ΔCT and ΔRpa49 (50 nM Rpa49CT) and 12 nM ΔRpa12. Percentage of fully extended and arrested transcripts in reference to the pulse labelled arrested transcript (time point 0) at the respective time points are depicted below each gel section.

**Suppl. Fig. 3.**
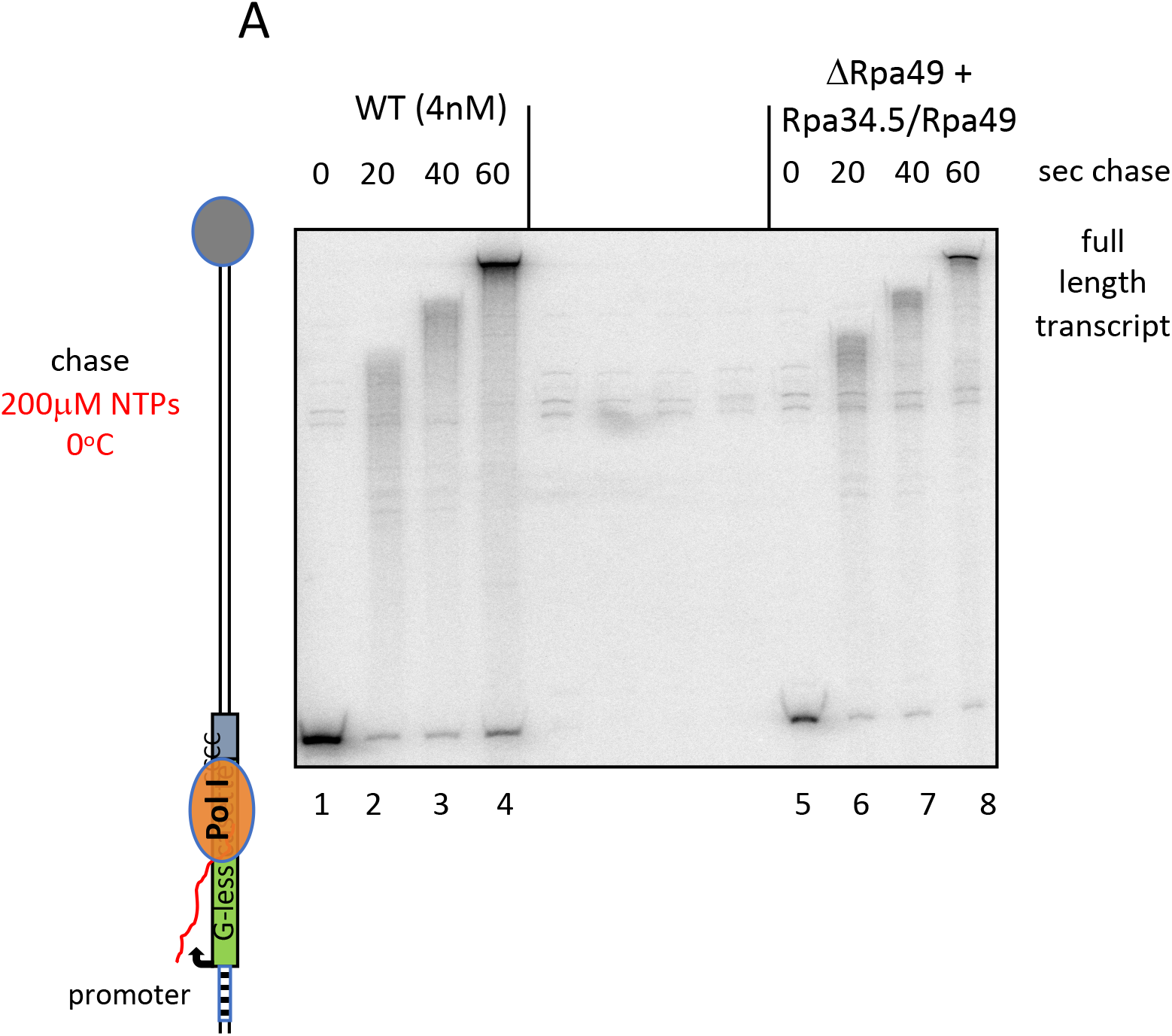
Original gel section of Fig 3A, lanes 1–8

**Suppl. Fig. 4.**
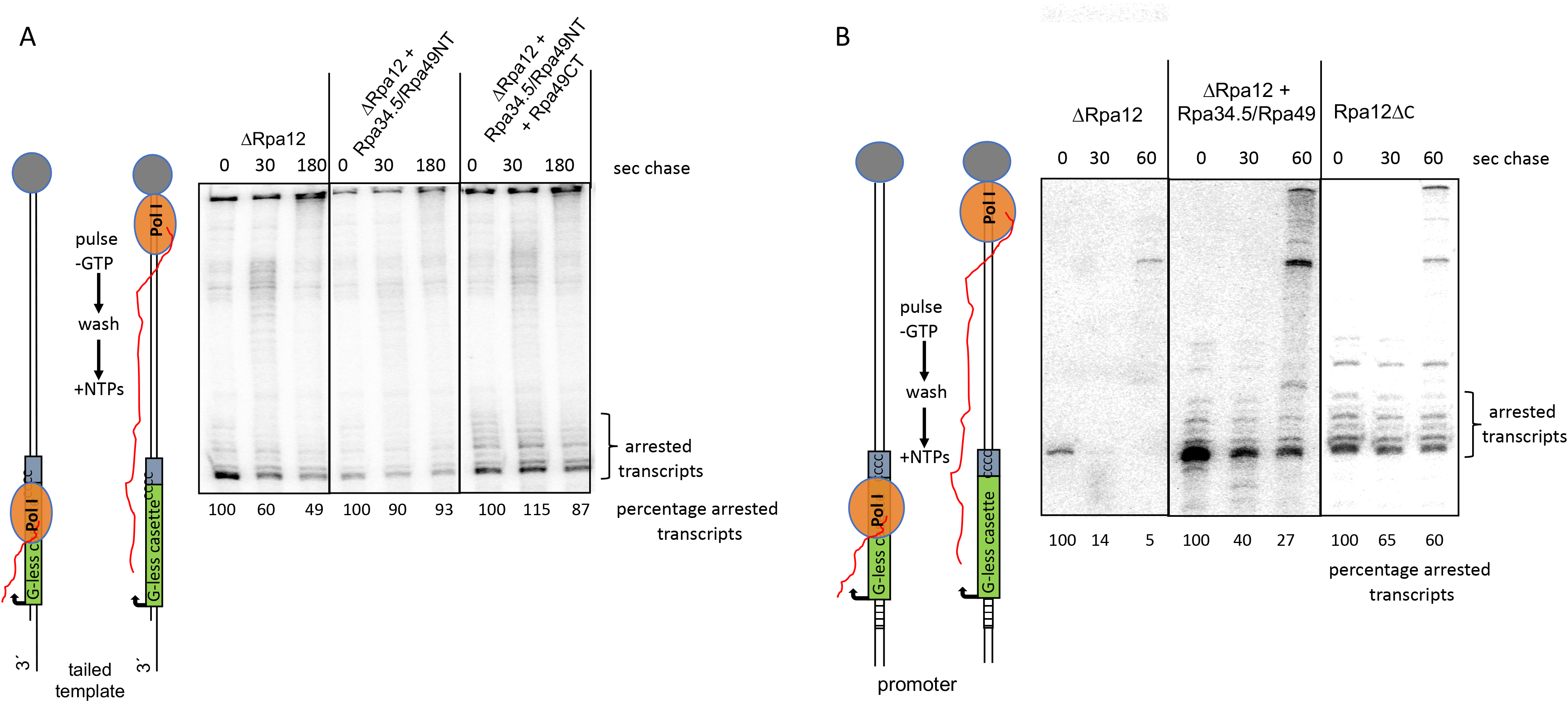
A) The dimerization domain without Rpa12.2 produces stalled transcripts at the G-less cassette which cannot be extended. Same experiment as in Fig. 5B, however using different batches of ΔRpa12.2 and heterodimer fractions. Concentration of heterodimer fractions was 25 nM instead of 50 nM. The strong band at the top of the gel is non-specific and comigrates with fully extended transcripts. Percentage of the arrested transcripts is shown below the single lanes. B) Production of non- extendable arrested transcripts at the G-less cassette depends on the presence of the heterodimer and the absence of Rpa12.2. Transcription complexes containing ^32^P-labelled RNA were generated using promoter-dependent transcription using the Pol I mutants and protein fractions as indicated and stalled at the G-less cassette (10 min pulse labelling without GTP). Complexes were washed twice, NTPs (200 μM end concentration) were added and the reactions were stopped after 30 and 180 seconds respectively. Transcripts were analyzed on a 6% polyacrylamide gel containing 6 M urea. Percentage of the arrested transcripts is shown below the single lanes.

**Suppl. Fig. 5.**
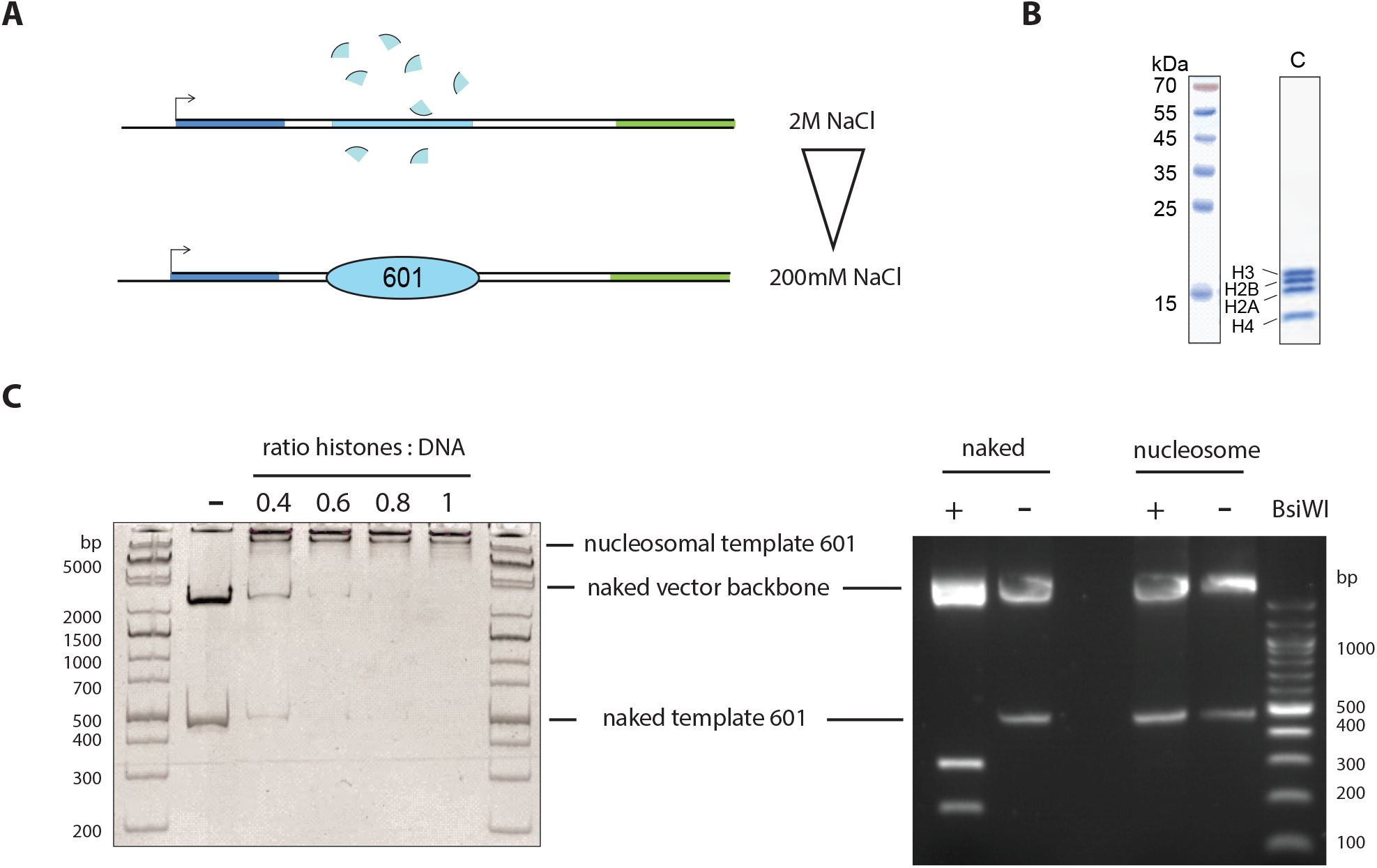
Purified core histones H2A, H2B, H3 and H4 from chicken erythrocytes were assembled according to (Rhodes and Laskey, 1989) using the salt gradient dialysis technique. A) Schematic outline. B) 12% PAGE (Coomassie stain) of purified chicken histones. C) A typical assembly reaction (50 μ.l) contained 5 μg DNA carrying the 601 nucleosome positioning sequence (Lowary and Widom, 1998), varying amounts of histone octamer, 200 ng BSA/ml, and 250 ng competitor DNA in high salt buffer (10 mM Tris, pH 7.6, 2 M NaCl, 1 mM EDTA, 0.05% NP-40, 2 mM p-mercaptoethanol). The salt was continuously reduced to 200 mM NaCl during 16–20 h. The quality of the assembly reaction was analysed on a 5% PAA gel in 0.4 x TBE followed by ethidium bromide staining. Different ratios of DNA to nucleosomes were analysed by electromobility shift assays to reveal optimal conditions for nucleosome assembly (left panel). Protection assay: 0.5 mg of naked and nucleosome associated template were incubated with and without restriction enzyme Nb.BsmI (NEB). Note that the 601 containing sequence was not cleaved in the presence of a nucleosome.

**Suppl. Fig 6.**
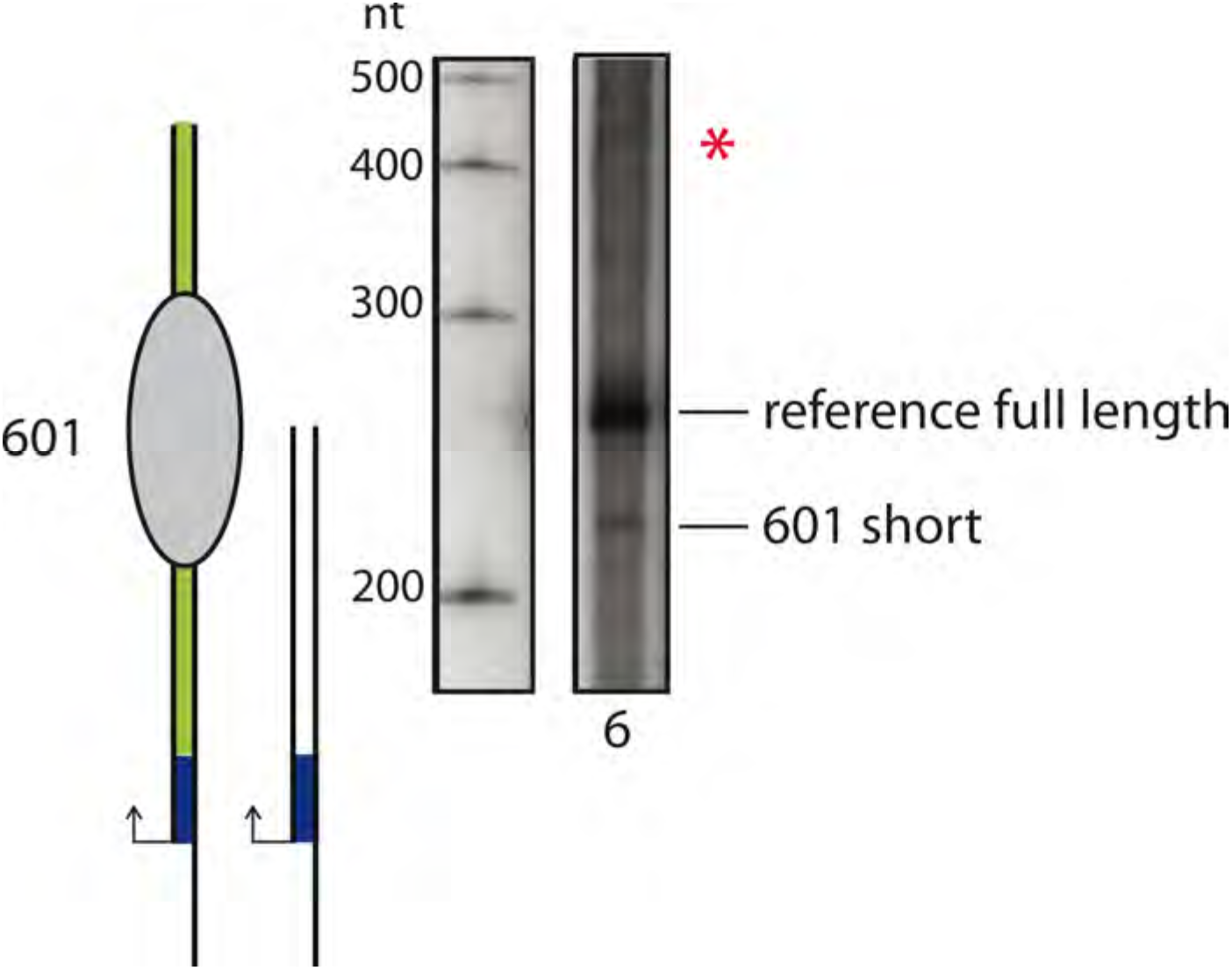
A shorter transcript is detected if Pol II transcription is arrested by a nucleosome on tailed templates. Longe exposure of lane 6 of Fig. 6 indicated an additional RNA which corresponded to the length of an arrested transcript.

## Supplementary

**Table 1:** Strains.

| Strain | Name | Parental strain | Genotype | Reference |
| --- | --- | --- | --- | --- |
| y2423 | BY4741 A135-TEV-ProtA | BY4741 | MATa, *his3-1 leu2-0 met15-0 ura3-0 rpa135::RPA135-TEV-ProtA-kanMX6* | (Hierlmeier et al., 2013) |
| Y2670 | yJPF162-1a | BY4741 | *his31; leu20; met15-0; ura3-0; RPA135-TEV-ProtA::kanMX6 HIS3MX::GAL::HA-RPA49* | (Pils et al., 2016) |
| Y2679 | yJPF165-1a | BY4741 | MATa, *his3-1, leu2-0, lys2-, ura3-0 RPA135-TEV-ProtA::kanMX6 HIS3MX::GAL::HA-RPA12* | (Merkl et al., 2014) |
| y2423 | BY4741 A135-TEV-ProtA | BY4741 | MATa, *his3-1 leu2-0 met15-0 ura3-0 rpa135::RPA135-TEV-ProtA-kanMX6* | (Hierlmeier et al., 2013) |
| y2424 | BY4741 Rpb2-TEV-ProtA | BY4741 | MATa, *his3-1 leu2-0 met15-0 ura3-0 rpb2::RPB2-TEV-ProtA-kanMX6* | (Hierlmeier et al., 2013) |
| y2425 | BY4741 C128-TEV-ProtA | BY4741 | MATa, *his3-1 leu2-0 met15-0 ura3-0 c128::C128-TEV-ProtA-kanMX6* | this study |

To obtain the yeast mutant expressing Rpa12.2ΔCT plasmid p1839 was transformed in strain y2679. Cultivation was in glucose-containing medium.

**Table 2:** Plasmids.

|  |  |  |  |
| --- | --- | --- | --- |
| 1247 | pUC19PIP g-601 | EcoRI/EcoRV digested amplicon from pRS316 SIRT G-less with 2107/2108 into KasI, blunted/EcoRI digested pUC19 601 | Merkl et al., 2014) |
| 1961 | pUC 19 PIP g- TER | contains Pol I promoter template with rDNA terminator including T1 T-rich element for in vitro transcription<br>SacI/BamHI cut terminator region amplified with primers #3050/3051 was cloned into SacI/BamHI cut vector #1247 | Merkl et al., 2014) |
| 2148 | pUC 19 PIP g- TER elongated | amplicon of primers #2479 and #2480 from pT11 phosphorylated and cloned into SacI cut #1961 | Merkel et al., 2014) |
| 2313 | pUC19 PIP g- elong 601 | extended g-less cassette to enhance labelling in the elongation assay; for promotor-dependent transcription | Merkel et al., 2014) |
| 435 | pMAX1 | SacI-fragment (3780 bp) from YepSIRT (Musters et al., 1989) containing ETS1 including Pol I promoter and 520 bp at the C-terminus of 25S rDNA cloned in pBluescript (6731 bp) | (Pilsel et al., 2016) |
| 1253 | pUC 19 tail G-601 | tailed template DNA containing 601 binding sequence for unspecific in vitro transcription | Merkel et al., 2014) |
| 1573 | pUC19 tail G- | tailed template DNA without 601 binding sequence for unspecific in vitro transcription | Merkel et al., 2014) |
| 2316 | pUC19 tail g- elong 601 | extended g-less cassette to enhance labelling in the elongation assay; for promotor-independent Trx | this study |
| 1839 | pCM182-LEU2-rpa12delC | plasmid expressing the N-terminus of Rpa12 (aa 1- aa 85) | this study |

**Table 3:**
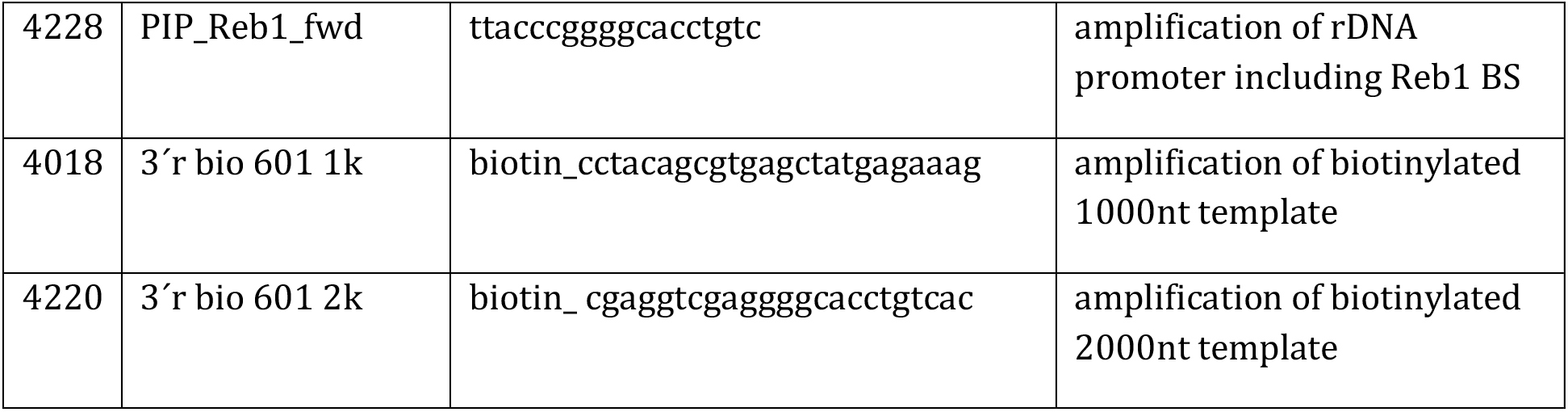
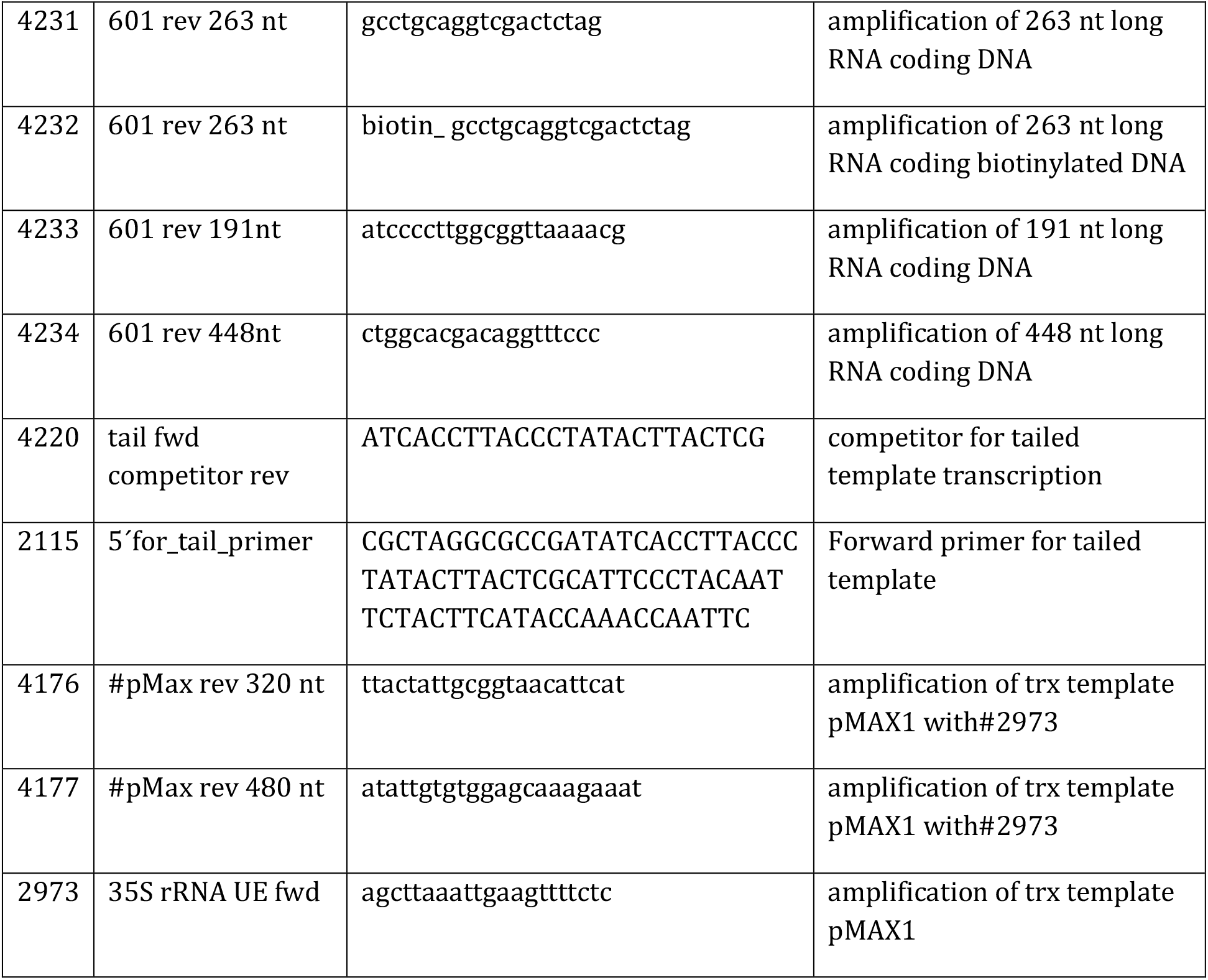
Oligonucleotides.

